# Deciphering shared molecular dysregulation across Parkinson’s Disease variants using a multi-modal network-based data integration and analysis

**DOI:** 10.1101/2024.09.30.615801

**Authors:** Alise Zagare, Irina Balaur, Adrien Rougny, Claudia Saraiva, Matthieu Gobin, Anna S. Monzel, Soumyabrata Ghosh, Venkata P. Satagopam, Jens C. Schwamborn

## Abstract

Parkinson’s disease (PD) is a progressive neurodegenerative disorder with no effective treatment. Advances in neuroscience and systems biomedicine now enable the use of complex patient-specific *in vitro* disease models and cutting-edge computational tools for data integration, enhancing our understanding of complex PD mechanisms. To explore common biomedical features across monogenic PD forms, we developed a knowledge graph (KG) by integrating previously published high-content imaging and RNA sequencing data of PD patient-specific midbrain organoids harbouring LRRK2-G2019S, SNCA triplication, GBA-N370S or MIRO1-R272Q mutations with publicly available biological data. Furthermore, we generated a single-cell RNA sequencing dataset of midbrain organoids derived from idiopathic PD patients (IPD) to stratify IPD patients towards genetic forms of PD. Despite high PD heterogeneity, we found that common transcriptomic dysregulation in monogenic PD forms is reflected in IPD glial cells. In addition, dysregulation in ROBO signalling might be involved in shared pathophysiology between monogenic PD and IPD cases.

## Introduction

The characteristic motor impairment in Parkinson’s disease (PD) is attributed to the gradual degeneration of dopaminergic neurons in the midbrain, yet the exact cause of this neuronal loss remains unknown^1^. Furthermore, genetics account for only approximately 10% of PD cases, leaving the majority of cases classified as idiopathic (IPD) ^2,3^. Identification of shared dysregulated molecular pathways between genetic and idiopathic PD cases holds significant importance in understanding disease mechanisms enabling the development of therapeutic strategies that could be applicable across multiple PD patient groups.

In recent years, there has been a significant advancement in high-throughput experimental technologies, allowing scientists to generate large amounts of biomedical data to investigate complex disease mechanisms^4^. However, this data is often collected and provided in various formats across different studies, hindering data integration and secondary analyses ^4,5^. Therefore,in systems biomedicine, harmonisation and standardisation of research outputs are crucial to maximise the interpretability and reproducibility of results, and to facilitate comprehensive data integration across various research studies and experiments. Graph databases (GDBs) have been used for this task in systems biomedicine due to their flexibility i) to represent naturally the biomedical information and to integrate large sets of heterogeneous data types (including omics, clinical, imaging, sensor data, etc.), ii) to capture complex data inter-relationships and iii) to provide support for network-based analysis and modelling of biomedical data^6–9^. This approach is particularly appealing in studying complex diseases, such as PD, where cause-effect relationships remain difficult to decipher ^10^. In particular, the integration of large amounts of heterogeneous data in knowledge graphs (KGs) enables the discovery of new relationships using reasoning frameworks ^11^ such as machine learning (ML)^12^, with various applications in biomedicine^13^. In the context of PD, several KGs have been developed for the identification of novel mechanisms and drug targets, either with a large biomedical scope only applied to PD^14–19^, or focusing on data related to neurodegenerative diseases^20–22^.

In this study, we developed a PD-related knowledge graph (PD-KG) integrating existing high-content imaging and RNA sequencing PD data with biological data from major public resources (e.g., Reactome^23^, IntAct^24^, DisGeNet^25^, DGIdb^26^, UniProtKB^27^). The integrated experimental data was previously acquired from midbrain organoids generated from induced pluripotent stem cells (iPSCs) of PD patients harbouring LRRK2-G2019S^28,29^, SNCA triplication^30^, GBA-N370S^31^ or MIRO1-R272Q^32^ mutations (Table M1). We further performed a network-based analysis on the PD-KG, focusing on the identification of common dysregulated molecular features across multiple PD-associated mutations. This analysis resulted in a comprehensive overview of the pathways, gene interaction partners and drugs shared between the datasets, revealing 25 genes with shared dysregulation in at least two monogenic PD cases. Notably, these 25 genes also demonstrated differential expression in the glial cells of idiopathic Parkinson’s disease (IPD) patients, as observed in a newly generated single-cell RNA sequencing experiment. Additionally, our analysis suggests that ROBO signalling may represent a potential shared disease mechanism between IPD and monogenic PD. Importantly, our work also provides a harmonised and integrated multimodal dataset comprising transcriptomics and high-content imaging data from monogenic PD and IPD-specific midbrain organoids, ready to use for future stud ies.

**Table M1.** Idiopathic PD patient and healthy control iPCS lines used to generate midbrain organoids for previously unpublished single-cell RNA sequencing and imaging datasets. IBBL - Integrated BioBank of Luxembourg; MPI - Max Planck Institute; WTSI – Wellcome Sanger Institute.

| Sample ID | Healthy/IPD | Age of onset | Age of sampling | Sex | Source |
| --- | --- | --- | --- | --- | --- |
| CTRL1 | Healthy | - | 65 | F | IBBL/MPI Muenster |
| CTRL2 | Healthy | - | 63 | F | IBBL/MPI Muenster |
| CTRL3 | Healthy | - | 68 | F | IBBL/MPI Muenster |
| CTRL4 | Healthy | - | 30 | M | Coriell (GM25256) |
| CTRL5 | Healthy | - | 55 | M | Coriell (GM23338) |
| CTRL6 | Healthy | - | 55-59 | M | Bill Skarnes, WTSI |
| IPD1 | IPD | 57 | 67 | F | IBBL/MPI Muenster |
| IPD2 | IPD | 62 | 67 | F | IBBL/MPI Muenster |
| IPD3 | IPD | 60 | 68 | F | IBBL/MPI Muenster |
| IPD4 | IPD | 64 | 68 | M | IBBL |
| IPD5 | IPD | 68 | 70 | M | IBBL |
| IPD6 | IPD | 59 | 60 | M | IBBL |

## Results

### PD Knowledge Graph (PD-KG)

We developed the PD-KG, integrating the high-content imaging and the top 100 significantly differentially expressed genes (DEGs) from RNA sequencing experiments available in the previously published experimental datasets on PD patient-specific midbrain organoids^28,30–32^ with biological data from public resources (e.g., Reactome^23^, IntAct^24^, DisGeNet^25^, Drug–Gene Interaction Database (DGIdb)^26^, UniProtKB^27^) corresponding to multiple biomedical layers (e.g. disease associations, drug targets, pathway involvements, protein-protein interactions). Concepts (such as genes/proteins, pathways, and drugs) were represented as nodes and relationships among concepts (e.g. protein-disease association, protein-drug target) as edges in the underlying graph. Annotations (such as the cell line provenience for imaging data, and the mapping between gene symbols and unique UniProt identifiers to enhance interoperability), were captured as attributes (properties) of nodes and edges in the graph. The PD-KG contains 3610 nodes and 8512 edges (relationships). Its data model is shown in Figure 1 and details on the semantics of the nodes and relationship types are provided in Supplementary Data. For example, the transcriptomics measurement of the TCEAL7 gene in the GBA_3 cell line at the Day 30 time point in the GBA-PD dataset is shown by the “GBA” edge (relationship) connecting the TCEAL7 gene and the “GBA_3 D30” CellLineTimePoint nodes (Figure2A).

**Figure 1.**
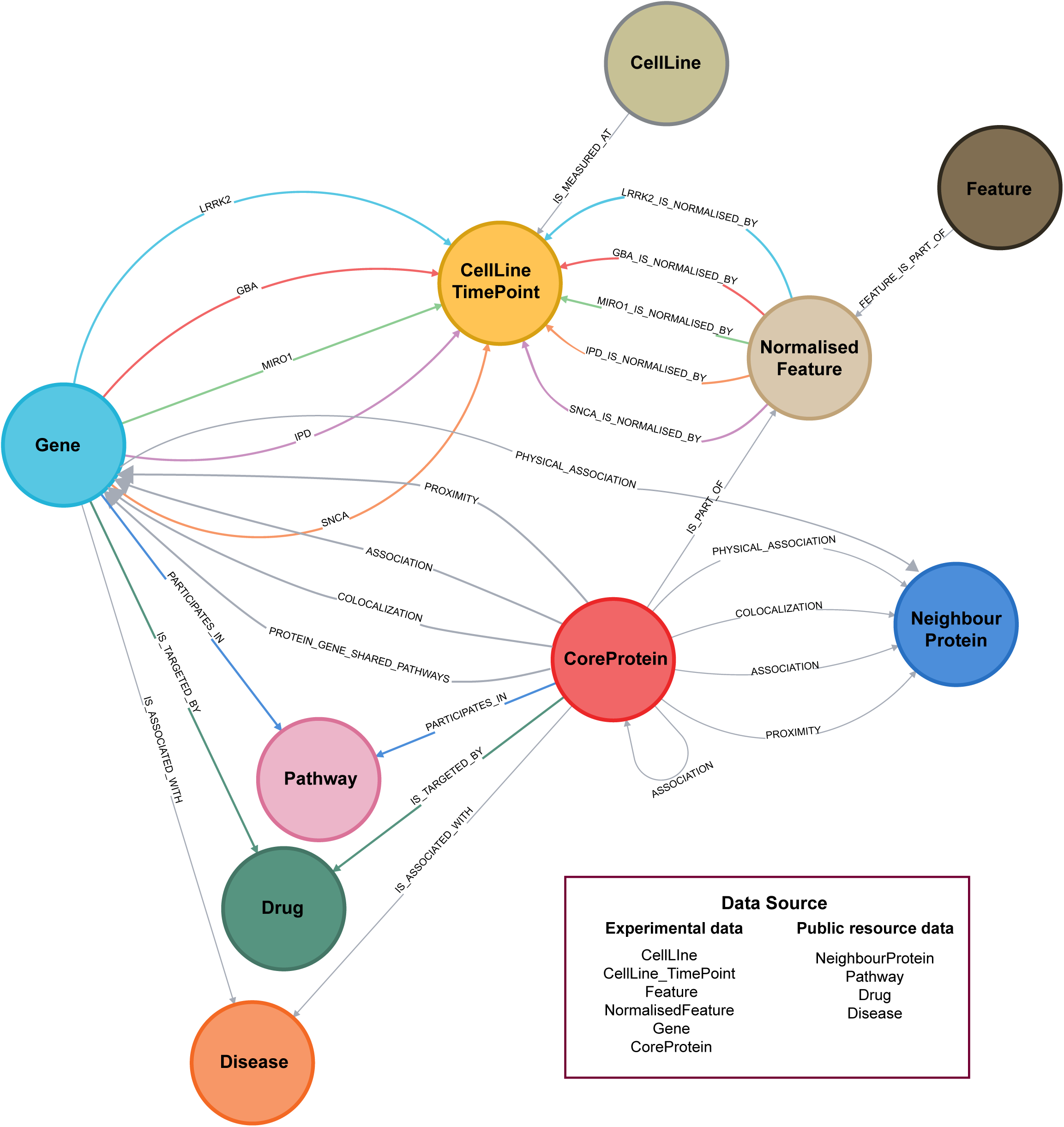
The graph data model for the PD-KG. The data types given by biological concepts such as proteins, genes, pathways, diseases, drugs, and cell lines are represented as nodes and their interrelationships (e.g. gene-pathway involvement) as edges (relationships) in the underlying graph.

### In-silico analysis reveals shared genes and molecular pathways between monogenic PD cases

Integration of data from previously published studies on PD patient-specific midbrain organoids allowed us to explore the similarities in gene and protein expression patterns between the four different PD forms caused by a single mutation in the LRRK2, SNCA, GBA or RHOT1 (encoding MIRO1) genes. We focused on the top 100 significant differentially expressed genes (DEGs) from each experimental dataset, which we believe adequately represent the key transcriptomic signatures underlying mutation-associated phenotypes, and we aggregated a combined set of 400 DEGs for all 4 studies. First, we noticed that there is no single shared DEG between all four monogenic PD cases. However, we observed 25 genes shared between at least two of the datasets (Figure 2A, Table S1). Interestingly, 15 of these 25 genes were shared between LRRK2-PD and MIRO1-PD datasets, suggesting a higher similarity of genetic dysregulation between these two monogenic PD forms. Furthermore, only 12 out of the total 25 shared genes showed the same expression direction (UP or DOWN) across the compared groups (Figure 2A - highlighted in bold, Table S2). The set of 13 genes with different regulation directions suggests that similar molecular processes may be differentially regulated in different PD cases.

**Figure 2.**
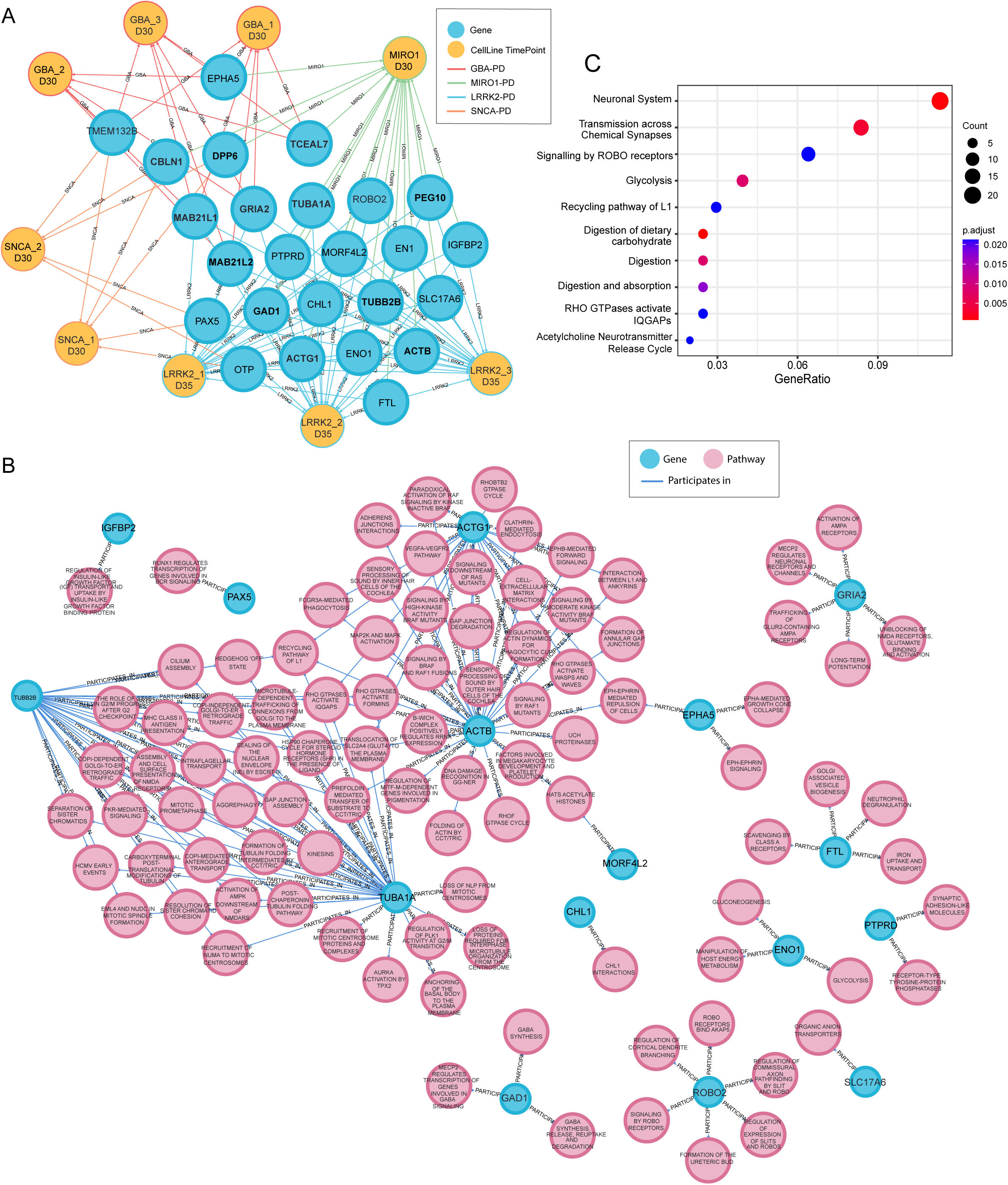
*In silico* analysis reveals shared genes and pathways between monogenic PD datasets. **A)** Visual representation of PD-KG subset demonstrating relationships between cell lines used for midbrain organoid generation in individual experiments (yellow nodes) and genes from the top 100 significantly differentially expressed lists (blue nodes). D30 and D35 indicate that transcriptomics analysis was done for midbrain organoids at day 30 or 35 of culture, respectively. The colour of the edges indicates different datasets (GBA-red, LRRK2- blue, MIRO1-green, SNCA-orange). **B)** Visual representation of PD-KG subset demonstrating shared genes between the four PD dataset involvement in pathways (pathway source: Reactome). Blue nodes – genes; pink nodes – pathways. **C)** Pathway overrepresentation analysis (ORA) of the merged list of the top 100 significantly differentially expressed genes from all four PD datasets.

Integration with public data repositories allowed us to further explore the experimental data. We used the PD-KG to identify common pathways between the 25 genes of interest shared between at least two datasets. Sixteen genes were reported as involved in at least one Reactome pathway (Figure 2B, Table S3). Genes MORF4L2, EPHA5, ACTB, ACTG1, TUBA1A and TUBB2B showed the highest involvement rate in a variety of cellular pathways. These six genes were associated with 69 pathways, including the “Recycling Pathway of L1”, “Translocation of SLC24 (GLUT4) to the plasma membrane”, “RHO GTPases activate IQGAPS”, “RHO GTPases activate FORMINS”, and “EPH-EPHRIN mediated repulsion of cells” (TableS3).

Further, we conducted an over-representation analysis (ORA)^33^, considering the combined list of the top 100 most significant DEGs from individual genetic experiments. While the PD-KG indicated specifically the Reactome pathways involving the 25 shared DEGs, the ORA analysis explored whether both the shared and unique DEGs from each study are involved in similar pathways. The ten most enriched pathways were associated with the neuronal system, synaptic function, ROBO signalling, metabolism (glycolysis, digestion of carbohydrates, digestion and absorption), IQ motif-containing GTPase-activating proteins (IQGAPs) and acetylcholine release cycle (Figure 2C). The overlap between several pathways associated with shared and experiment-specific transcriptomic features suggests commonalities in pathway-level dysregulation among the four PD mutations.

The available high-content imaging data were inconsistent between the initial four midbrain organoid experiments. This was due to the custom selection of proteins for imaging analysis and the difference in the relevant time points considered in each individual experiment. The proteins for immunostainings analysed in the individual studies were always selected based on previous knowledge of predicted mutation-associated phenotypes^28,30–32^. Across all four independent experiments, a set of 12 core proteins was analysed (Table S4). The tyrosine hydroxylase (TH, UNIPROT id: P07101) was the only core protein common to all datasets. TH is a rate-limiting enzyme in dopamine synthesis and, thus, an essential marker of dopaminergic neurons, which is the main neuronal population affected in PD. Given the limitations that hinder the core protein abundance comparison across all four datasets, we used the PD-KG and explored the pathways involving both the core proteins from imaging data and the top 100 significant DEGs within the same experiment, revealing genotype-phenotype relationships and the relevance of the analysed proteins (Figure S1A-C, Table S5). We observed that in the LRRK2-PD experiment, there were no connections between the top DEGs and the core proteins. The other three datasets shared several core proteins, namely TUJ1, TH, GFAP, and S100b, and we focused on their pathway involvement, as follows. In the MIRO1 and SNCA datasets, the GFAP and S100b glial markers from imaging data shared pathways with the cytoskeleton filament-associated STMN1and GFAPgenes, suggestingthat glial cell development or maturity might be affected in these forms of PD (Figure S2A-B). In the GBA study, relationships were identified between the TH and SOX2 proteins and one or several significant DEGs. This indicates that dysregulation in stem cells and dopaminergic neurons plays a crucial role, particularly in the development of GBA-PD as also reported in the original study^31^ (Figure S2C). In all three datasets, the TUJ1 protein shared pathways with several significant DEGs, including CDKN1A, ACTB, VASH2, L1CAM, CTSF, PMSD5, and others (Figure S2A-C). TUJ1 is a neuronal protein that forms microtubules and is involved in neurogenesis and axon guidance. Overall, associations between glial proteins and neuronal TUJ1 with the significant DEGs involved in cytoskeleton dynamics are consistent with ORA results, suggesting common PD dysregulation in ROBO signalling, also involved in axon guidance and cytoskeleton organisation^34^. Although, in the LRRK2 dataset we did not find shared pathways between analysed core proteins and the top significant DEGs (TUJ1 was not in the imaging dataset), several isoforms of tubulins, such as TUBA1A, TUBB2B and TUBA1B were found in the list of top 100 significant DEGs, further suggesting that disrupted microtubule cytoskeleton organisation and altered axon guidance might be shared mechanisms between the four genetic PD cases.

### Idiopathic PD shares transcriptomic dysregulation with monogenic PD cases

The integration of transcriptomic data of four different PD-associated mutations in the PD-KG revealed 25 significantly dysregulated genes shared between at least two of the mutations. Moreover, we were able to identify dysregulation in tubulins and ROBO signalling indicating cytoskeleton organisation and axon guidance as a potential common PD mechanism across different PD-associated mutations. This suggests that despite the overwhelming PD heterogeneity, there are some similarities in the transcriptomic landscape between LRRK2-G2019S, 3xSNCA, GBA-N370S and MIRO1-R272Q genetic PD forms. We further investigated if there are also similarities between these four genetic PD cases and IPD.

We performed single-cell RNA sequencing on iPCS-derived midbrain organoids from six (three female and three male) IPD patients and six (three female and three male) healthy controls (Table M1). Using cell type-specific markers^35,36^, we identified nine distinct cellular populations - GABAergic neurons, subdivided into mature and young GABAergic neurons, dopaminergic neurons, subdivided into mixed and vulnerable populations, neuroblasts, radial glia, neuronal stem cells, astrocytes and oligodendrocytes (Figure 3A, Figure S2) that were all present in IPD and healthy control samples (Figure 3B). The two dopaminergic neuron clusters were named vulnerable and mixed according to the expression of vulnerability markers ^37^ (Figure S3).

**Figure 3.**
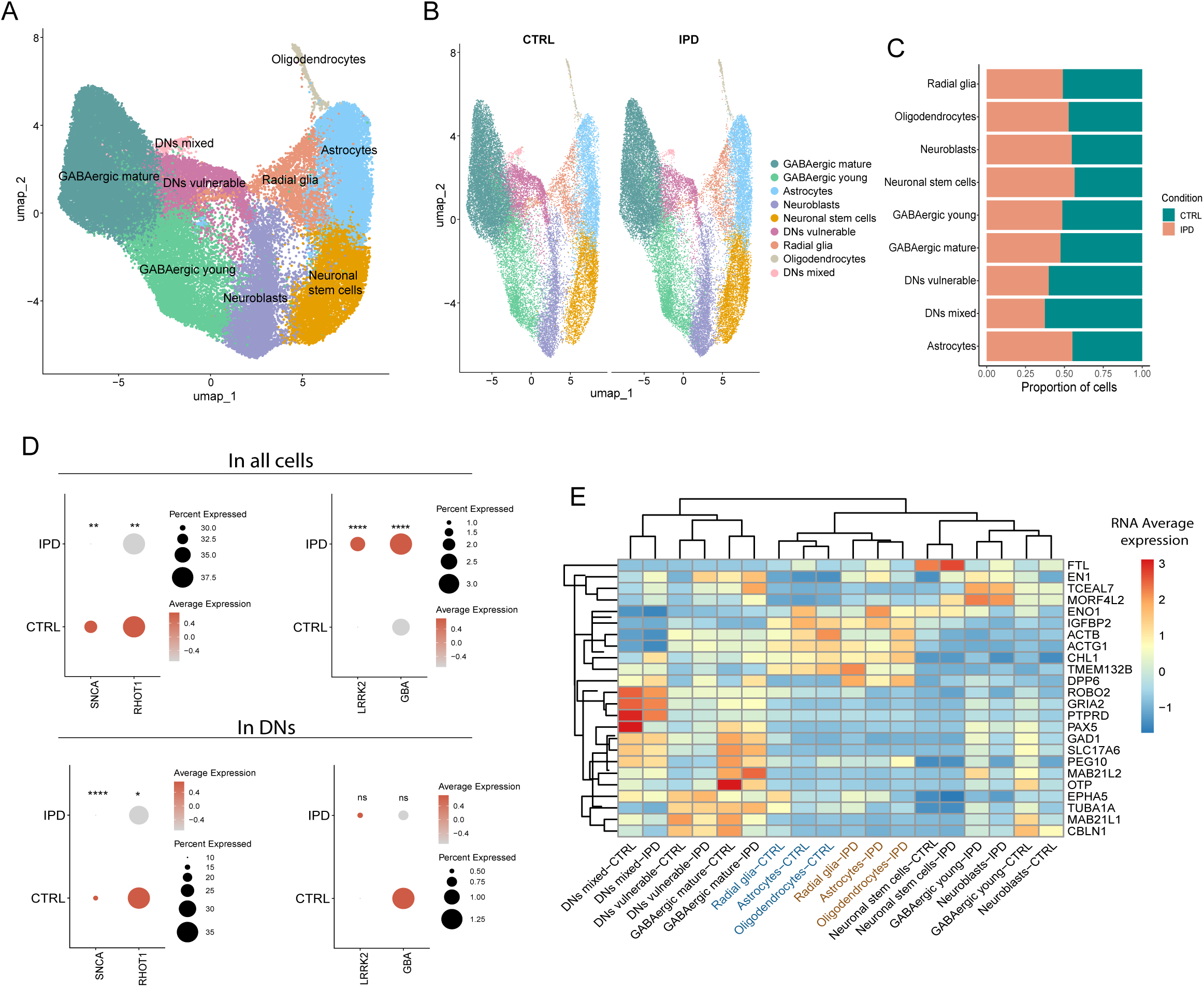
Single-cell RNA sequencing analysis of IPD-patient-specific midbrain organoids. **A)** UMAP representation of identified cellular populations. Every dot represents a single cell and colours indicate different cellular populations. **B)** UMAP representations of cellular populations separated by healthy control (CTRL) and IPD samples. Every dot represents a single cell and colours indicate different cellular populations. **C)** Proportion of each cell population in CTRL and IPD conditions. **D)** Scaled expression of SNCA, RHOT1, LRRK2 and GBA genes in IPD and CTRL samples considering the expression across all cells or a subset of dopaminergic neuron clusters (DNs mixed and DNs vulnerable). Dot size represents the percentage of cells expressing the gene, colour represents the gene expression level. Statistical significance was determined using the Wilcoxon rank-sum test. **E)** Unsupervised clustering of cell populations within IPD and CTRL samples based on the average cell expression of the 25 genes of interest shared between monogenic PD datasets.

A significant reduction in TH-positive dopaminergic neuron levels was reported in the four datasets of midbrain organoids derived from monogenic PD patient iPSCs, regardless of the mutations carried by the patients^28,30–32^. Thus, first, we wanted to confirm that IPD midbrain organoids also show a reduction of dopaminergic neurons compared to healthy controls. We assessed the percentage of each cell type in control and IPD conditions (Figure 3C, Figure S4) and observed that in the IPD case, there is a 1.7 times reduction of mixed dopaminergic neuron population and 1.5 times less vulnerable dopaminergic neurons. Additionally, GABAergic neuron populations and radial glia cells were reduced in IPD compared to control samples. However, the astrocyte, oligodendrocyte, neuroblast and neuronal stem cell populations were slightly larger in IPD than in healthy control midbrain organoids (Figure 3C, Figure S4).

Further, we checked the expression pattern of PD-associated genes (LRRK2, SNCA, GBA and RHOT1 encoding MIRO1) corresponding to the monogenic PD datasets. We observed that SNCA and RHOT1 expression levels were significantly reduced in all cells as well as in dopaminergic neurons of IPD samples compared to healthy controls (Figure 3D). LRRK2 expression was significantly higher in IPD samples taking all cells together but in dopaminergic neurons the expression difference was insignificant. Similarly, GBA was significantly more expressed in IPD samples analysing its bulk expression; however, it showed lower expression in IPD dopaminergic neurons compared to healthy control samples. The significant dysregulation of genes associated with monogenic forms of PD suggests their potential role in IPD development and shared molecular disease mechanisms between IPD and monogenic PD.

To investigate if there is a similar genetic dysregulation between PD monogenic forms and IPD, we inspected the expression level of the 25 genes of interest shared between at least two monogenic PD experiments (Figure 2A, Table S1) in the IPD single -cell RNA sequencing dataset. We observed that the expression pattern of these 25 genes separated the IPD and the healthy control glial populations - radial glia, astrocytes and oligodendrocytes (Figure 3E). Moreover, control glial cells were clustered together with neuronal cell types, while IPD glial populations were closer to stem cell, neuroblast and young neuron cellular populations, suggesting a reduced maturity state of IPD glial cells.

Finally, we determined the significant DEGs between IPD and healthy control midbrain organoids (bulk-mode). In total, there were 6321 significant DEGs (p.adjust<0.05). ORA analysis showed that the top 100 significant DEGs between IPD and control samples are involved in platelet functions, metabolic processes, and regulation of SLIT/ROBO signalling involved in axon guidance (Figure 4A). The occurrence of ROBO signalling among the most enriched pathways based on the top 100 DEGs in IPD, suggests that ROBO signalling might play an important role in PD, independently of disease aetiology, providing a potential link between the pathophysiology of genetic and idiopathic PD. Next, we explored the genes associated with ROBO signalling in individual datasets (Figure 4B). A set of 18 genes across four monogenic PD datasets and the IPD dataset showed involvement in ROBO signalling-associated pathways as follows LRRK2 (7 genes), SNCA (2 genes), GBA (one gene), MIRO1 (3 genes) and IPD (6 genes).

**Figure 4.**
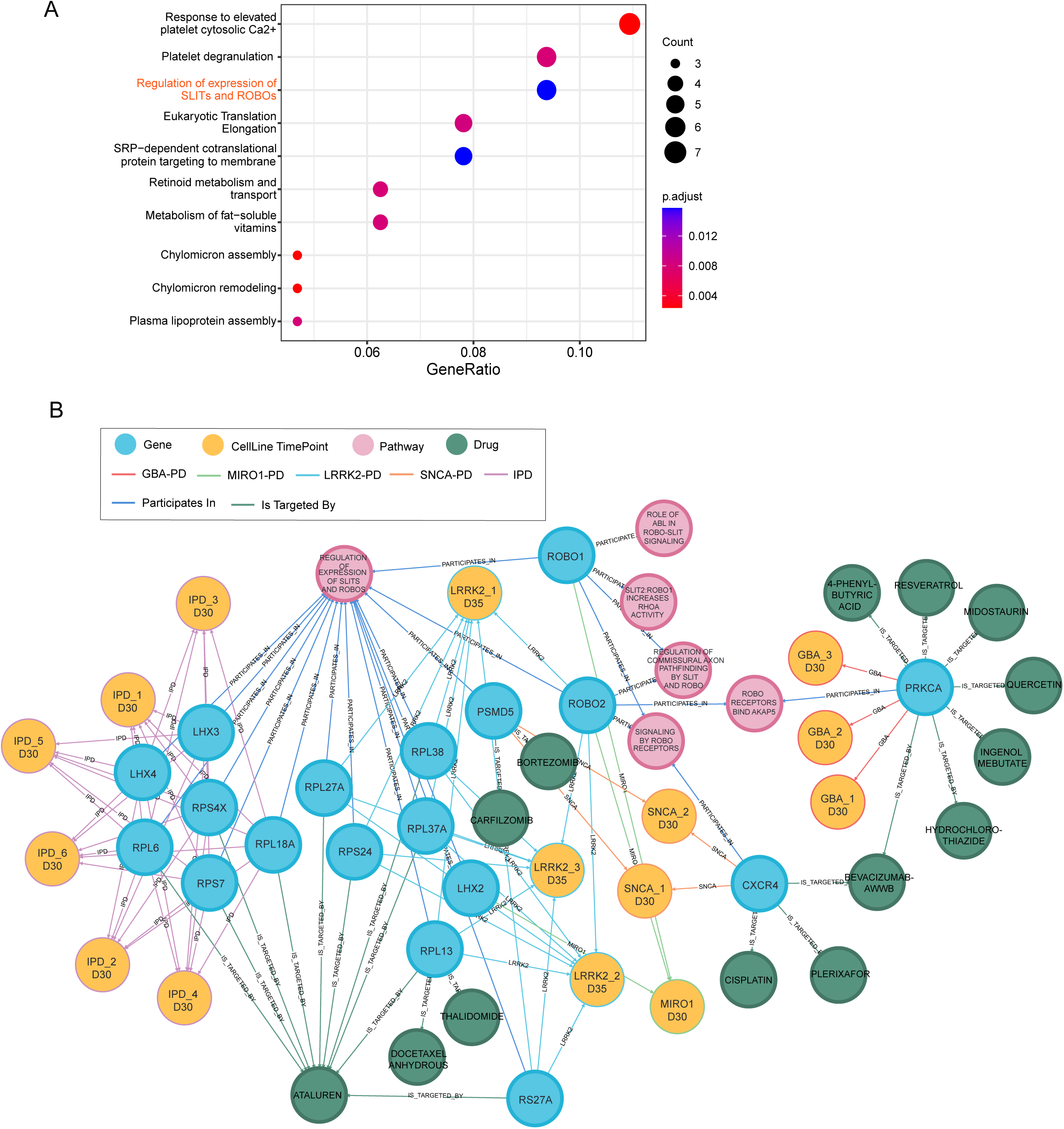
ROBO signalling dysregulation in monogenic and idiopathic PD. **A)** Pathway overrepresentation analysis (ORA) of the top 100 significantly differentially expressed genes between IPD and CTRL samples. Average gene expression across all cells considered for differential expression analysis. ROBO signalling as a re-occurring pathway is highlighted in red. **B)** Visual representation of PD-KG subset demonstrating relationships between cell lines (yellow nodes) and genes (blue nodes) involved in pathways associated with ROBO signalling (pink nodes) from four genetic PD and IPD datasets of the top 100 significantly differentially expressed genes with drugs from DGIdb (green nodes). The colour of the edges indicates different datasets (GBA-red, LRRK2-blue, MIRO1-green, SNCA-orange IPD-purple).

We then used the PD-KG to identify the drugs targeting genes involved in the ROBO signalling-associated pathways across all datasets. We determined a set of 14 such drugs, including medications used in cancer treatment (Thalidomide, Plerixafor, Midostaurin, Docetaxel Anhydrous, Cisplatin, Carfilzomib, Bortezomib, Bevacizumab -AwwB), treatment of metabolic disorders (4-Phenylbutyric acid) as well diuretic medications (Hydrochlorothiazide), plant-derived chemical compounds with pleiotropic effects (Resveratrol, Quercetin, Ingenol mebutate) and medication used for the treatment of Duchenne muscular dystrophy (Ataluren) (Figure 4B, Table S7). The list of drugs approved for other diseases that target the same genes found to be significantly dysregulated in PD may serve as a data-driven hypothesis for future PD treatment studies.

### Stratification of Idiopathic PD

We observed that LRRK2, SNCA, GBA and RHOT1 genes are significantly differentially expressed in IPD midbrain organoids compared to healthy controls. This suggests that at least some IPD samples should exhibit similar phenotypes to those associated with a mutation in the respective genes. To see if the IPD transcriptomic signature is similar to monogenic PD, we performed principal component analysis (PCA) and unsupervised hierarchical clustering on the top 100 significant DEGs for each mutation and IPD against healthy controls. Considering the technical variability of sequencing data (bulk or single -cell), for the stratification analysis we used log2 fold change PD vs control, calculated for each dataset separately. The PCA displayed the high heterogeneity of PD, positioning almost every monogenic PD case in a separate quadrant of the PCA plot (Figure 5A). The IPD and LRRK2-PD were clustered closer to the zero axis, suggesting more similar disease mechanisms. Similarly, in the heatmap of unsupervised hierarchical clustering, we observed that IPD and LRRK2-PD were clustered together, while other PD forms showed distinct transcriptomic signatures (Figure 5B). When looking at IPD cases individually, we observed that the first two data dimensions explain only about 50% of the variance with the most separation across the first dimension, highlighting the PD complexity (Figure 5C). In the heatmap, we observed that most IPD samples were rather clustered together, with the LRRK2 -PD and MIRO1-PD, except IPD2 with a transcriptomic signature more similar to GBA-PD (Figure 5D).

**Figure 5.**
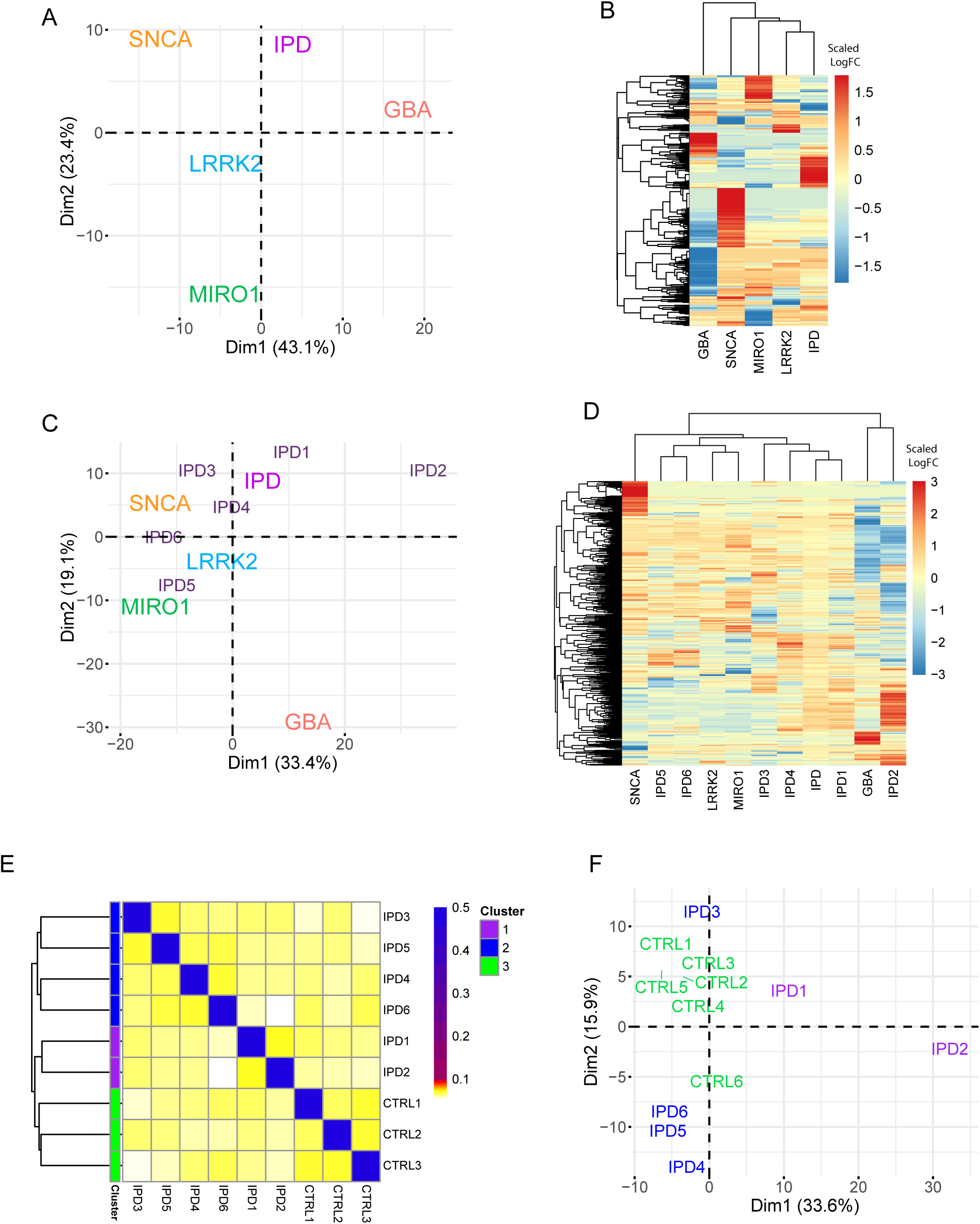
IPD stratification. **A)** Two-dimensional PCA plot of the monogenic PD and IPD datasets. **B)** Unsupervised clusteringof monogenic PD and IPD datasets. **C)** Two-dimensional PCA plots of the monogenic PD datasets and IPD datasets, considering every individual IPD sample separately. **D)** Unsupervised clustering of monogenic PD and IPD datasets, considering every individual IPD sample separately. For panels, A-D log2 FC of individual experiments considered (PDvsCTRL) of the genes included in the merged list of the top 100 significantly differentially expressed from each individual experiment. **E)** Similarity network fusion analysis (SNF) on integrated imaging and transcriptomic features (top 100 significantly differentially expressed genes) from 6 IPD and 3 CTRL cell lines. The computed similarity score among different cell lines is within the [0, 0.1] range in this set. The self -similarity has a value of 0.5. The colour palette shows the similarity among the cell lines: “white” for values close to 0, “yellow” for low values within the range [0, 0.1], “red” for values closer to 0.1, and “blue” for self -similarity (0.5). **F)** Two-dimensional PCA plot of IPD and CTRL samples, considering the average cell expression of the top 100 significantly differentially expressed genes between every IPD line against all six CTRL cell lines used for single -cell RNA sequencing.

In addition, we performed a stratification analysis for the individual IPD-patient-specific midbrain organoid cell lines against healthy controls using a similarity fusion network (SNF) approach^38^. We integrated the set of the top 100 significant DEGs and the imaging data of IPD midbrain organoids comprising quantification of 8 core proteins (namely TUJ1, S100B, GFAP, TH, MAP2, PAX6, KI67, SOX2), and TH fragmentation index from 6 IPD cell lines and the 3 healthy control cell lines (Figure 5E). Of note, here we selected only the 3 healthy control cell lines that were included in both imaging and transcriptomics data as the SNF method requires that the sample set is consistent across the data types layers (all samples are described by all considered data type layers)^38^. The predefined number three of SFN clusters separated control samples from IPD samples, subdividing IPD samples into two groups. One of the IPD clusters included three male (IPD4, IPD5 and IPD6) and one female (IPD3) IPD patient-derived samples (Figure 5E cluster 2). The second IPD cluster included the other two female (IPD1 and IPD2) IPD patient-specific samples (Figure 5E cluster 1), suggesting rather sex-independent PD mechanisms. Furthermore, the PCA approach on the transcriptomic signature of the top significant DEGs, including all six control lines, supported the clustering of IPD1 and IPD2 lines by SNF analysis (Figure 5F). The same IPD1 and IPD2 samples were also separated from other IPD lines and controls in Figure 5D; IPD2 clustered with GBA-PD and IPD1 being the farthest away from the LRRK2 and MIRO1 clusters. Clustering of IPD3, IPD4, IPD5 and IPD6 closer to LRRK2-PD and MIRO1-PD (Figure 5C-D), and at the same closer to healthy controls (Figure 5F) suggests that they might share disease mechanisms with the LRRK2-G2019S and MIRO1-R272Q associated PD. In addition, it also suggests that the disease course of these patients might be milder since the transcriptomic signature is more similar to all six healthy controls (Figure 5F).

Finally, we ran a core analysis on gene expression in the Ingenuity Pathway Analysis (IPA) platform^39^ to predict the significantly enriched pathways and their activity level for the individual IPD cases (Figure S5). The analysis revealed the molecular heterogeneity of IPD demonstrating a variety of dysregulated metabolic and signalling pathways, including ROBO signalling. The predicted activity (activation or inhibition) based on the log2 fold change values often showed an opposite trend for different IPD samples regarding the same pathway; for example, this was the case for protein ubiquitination, synaptogenesis, and mitochondrial dysfunction, again suggesting that the same molecular processes may be differentially regulated in different IPD cases.

Altogether these results show that IPD can be stratified toward genetic cases, which is important for targeted treatment approaches and a step towards personalised medicine.

## Discussion

Contextualisation using already-established knowledge is an important and often forgotten tool that could facilitate biomedical result interpretation. Moreover, secondary data analysis is important in order to make use of previously generated data as such, enhancing the potential of every dataset. Here, we demonstrated an example of experimental data integration with external data sources, enabling better data contextualisation and exploration using previous knowledge. The main limitation was that the available imaging datasets did not share most of the quantified features (core proteins), and image acquisition was done at different time points for different organoid models. However, identification of these limitations will allow us to further develop and optimise our experimental designs and expand the network by including missing imaging data. Moreover, in the current study, for the analysis, we considered only imaging data acquired at the closest time point to transcriptomics data generated only for one of the time points of organoid culture. In the future, the PD-KG can be complemented with transcriptomics data of other time points to explore genotype -phenotype relationships over time. Despite these limitations, we were able to find common disease -associated features and tendencies between monogenic and idiopathic forms of PD. In addition, the integration of external data sources into PD-KG allowed the exploration of shared dysregulated pathways and approved drugs targeting dysregulated genes, which indicates the benefits of using network-based approaches for biomedical data exploration, analysis and visualisation, and hypothesis derivation. Additionally, in our lab, the PD-KG can aid in designing future experiments by comparing core protein expression across different healthy control cell lines at various time points. This approach will help in selecting the most robust cell lines for comparisons with PD-patient cell lines.

Although we did not identify a single DEG shared among the LRRK2 -G2019S, 3xSNCA, GBA-N370S and MIRO1-R272Q datasets, highlighting the heterogeneity of PD, we did identify 25 genes commonly dysregulated in at least two of these datasets. Furthermore, these 25 genes were seen as involved in pathways closely linked to those associated with the unique genes from all four datasets, including IQ motif-containing GTPase-activating proteins (IQGAPs), glycolysis and translocation of glucose transporter GLUT4, as well as pathways involved in the organisation of the cytoskeleton, axon guidance and neuronal migration, such as ROBO signalling and recycling pathway of L1 and pathways regulating neuronal connections and communication such as EPH-ephrin mediated repulsion of cells and synaptic transmission. These results indicate that distinct PD transcriptomic signatures might still lead to dysregulation of the same molecular mechanisms.

Moreover, ROBO signalling was also among the top most enriched pathways of the significant DEGs between IPD and healthy controls. ROBO signalling regulates neuron migration, and axonal guidance, and is also involved in the control of the balance between cell proliferation and differentiation^34^. Studies in mice show that ROBO signalling is involved in the regulation of dopaminergic neuron projections^40^. In addition, dopaminergic neurons derived from human embryonic cells show ROBO2 protein expression increase over time and display robust response to axonal guidance cues by SLIT2, which is regulated by ROBO2 ^41^. Additionally, downstream effectsof the SLIT/ROBO signalling are associated with microtubule cytoskeleton organisation, including GTPase regulation^42^. While MIRO1 protein is a GTPase, LRRK2 is known to regulate small GTPases, which are regulators of actin cytoskeleton dynamics ^43–45^. In addition, cytoskeleton filaments actin (ACTB) and different subtypes of tubulins (TUBB3, TUBA1A, TUBB2B) were among the top 100 significant DEGs in the LRRK2 and MIRO1 PD datasets. The common dysregulation of cytoskeleton dynamics might link the LRRK2 -G2019S and MIRO-R272Q-associated PD. Accordingly, the majority of the 25 shared DEGs were between these two PD datasets and they were positioned closer in the clustering analysis. However, in all four genetic PD datasets and also in IPD dataset, we could identify significant DEGs associated with the SLIT-ROBO pathway (Figure 4B), suggesting that SLIT/ROBO might play an important role in PD independently of disease aetiology, providing a potential link between the pathophysiology of genetic and idiopathic PD. In the PD-KG we could see that multiple of the genes involved in the regulation of SLIT proteins were ribosomal, indicating potential aberrant protein translation affecting SLIT protein functionality. Interestingly, the drug, namely Ataluren, targeting 4 out of the 6 IPD genes linked to SLIT proteins: RPL18A, RPL6, RPS7 and RPS4X, acts by interacting with the ribosomes, and recruiting near-cognate transfer-RNAs to ensure the full-length-protein synthesis frommessenger-RNA, surpassingthe nonsense mutations^46^. Missense mutations account for a large proportion of human diseases, and currently, RNA targeting therapies are among the successful approaches for treating genetic disorders caused by missense mutations^47^. It is known that PD is a multifactorial disorder and genetics together with environment and age play a role in the disease pathophysiology^3^. Thus, it is feasible that there are missense mutations that contribute to the development of PD cases classified as idiopathic. Additionally, the PD-KG allowed us to identify several other existing medications targeting dysregulated, SLIT/ROBO signalling - associated genes across all included datasets. Importantly, drug-gene network analysis can be also expanded to other dysregulated genes, highlighting the practical value of the PD-KG in data contextualisation and the generation of new hypotheses, which could be valuable for future PD drug screening studies.

Interestingly, our analysis revealed that radial glia, astrocyte, and oligodendrocyte populations in IPD samples clustered with a less mature neuronal population (neuronal stem cells, neuroblasts and young GABAergic neurons) based on 25 genes with shared dysregulation between monogenic PD cases. This suggests that some glial properties or functions could be compromised in IPD, leading them to resemble a less mature cellular state. Glial cells are known to provide metabolic support to neurons. It has been suggested that in neurodegenerative diseases, glial cells undergo metabolic changes that enhance neuroinflammatory responses while reducing their neuroprotective and supportive functions^48^. Accordingly, ORA indicated that several metabolism-related pathways were among the enriched ones based on the top 100 significant DEG set. These metabolic alterations may be a result of the inflammatory phenotype driving glial metabolic dysfunction, which in turn contributes to neuronal metabolic changes. Further investigation of the expression of these 25 genes at the single-cell level in monogenic PD is needed to determine whether the altered glial cell transcriptomic profile represents a common mechanism in PD.

Aiming to stratify IPD, we extended the dataexploration and analysis fromthe PD-KG by using a set of methods and tools (including ORA, unsupervised hierarchical clustering, PCA and SNF) that provided complementary details on the upper biological levels (enriched pathways or biological processes and functions) rather than on single molecules (dysregulated protein or gene) across the input datasets. Moreover, the results from these analyses were obtained from quantitative information (such as transcriptomics or imaging data from the IPD dataset, log2 fold change between IPD samples and healthy controls) rather than on descriptive information integrated into the PD-KG (such as protein-pathway involvement). First, the ORA results on the enriched pathways for the genetic and IPD datasets were in agreement with the PD-KG results on the pathways shared across molecules of interest. The SNF analysis, which focused on exploring the similarities and differences within the IPD dataset by integrating the two layers of data available, specifically the cell-line-specific imaging data and the transcriptomics, indicated a separation between the control versus IPD samples. The SNF analysis also anticipated subtype identification across the IPD cell lines, which we also could observe in the PCA. Although stratification of IPD holds a great promise in the context of personalised medicine and targeted clinical studies, our main limitation was the number of cell lines available as well as the relatively low number of features considered within each data layer, especially in the imaging data.

The current version of the PD-KG can be enhanced through the integration of additional types of data and their relationships, including experimental data such as metabolomics and proteomics, as well as open-source resources such as the Human Protein Atlas^49^. This integration would increase the relevance and robustness of the generated hypotheses. The PD-KG can be also enriched with midbrain organoid data from PD patients carrying other PD-associated mutations. Given that midbrain organoids rather represent a developing brain at embryonic stages and therefore primarily reflect early disease mechanisms, additional integration of published data from PD patients would be of great importance. This integration would not only help to validate whether midbrain organoids can accurately model PD molecular mechanisms but also enable further exploration of shared molecular mechanisms across different forms of PD, particularly those influenced by ageing.

In conclusion, our study presents a comprehensive multimodal data integration and analysis approach for PD organoids (Figure 6). We curated and normalised a robust dataset encompassing transcriptomics and high-content imaging data from both genetic and idiopathic PD organoid studies. We also developed the PD-KG, a knowledge graph that integrates diverse biological data relationships from well-established public repositories, enhancing the exploration and contextualisation of PD experimental data. We used these components for IPD stratification, performing several complementary network-based and enrichment pathway analyses, which allowed a more detailed exploration of the experimental datasets. Importantly, using the created PD-KG we were able to derive a hypothesis of altered cytoskeleton dynamics and dysregulated ROBO signalling as common disease mechanisms between the genetic PD forms included in this study, which was further confirmed in the newly generated IPD single-cell RNA sequencing dataset.

**Figure 6.**
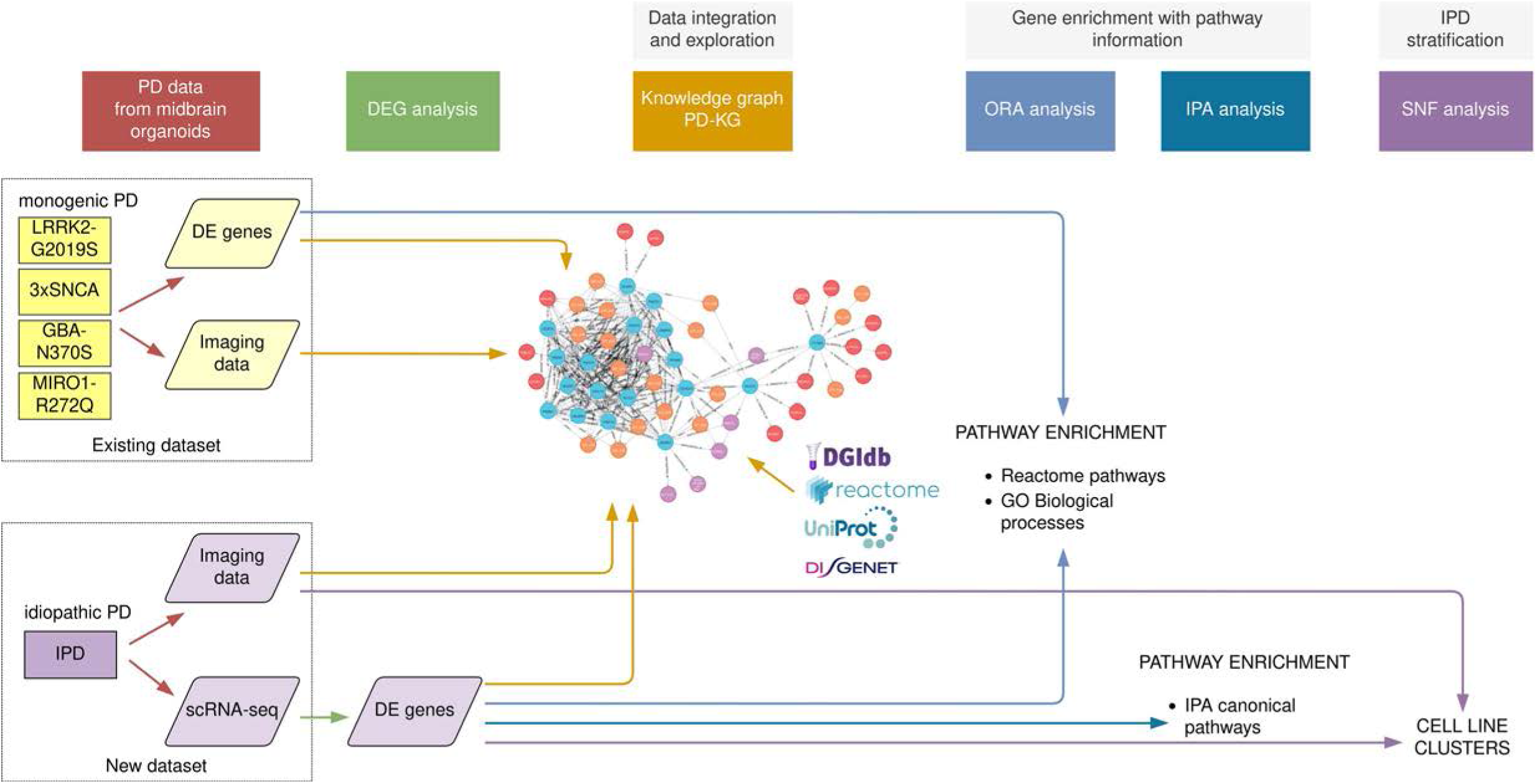
**Schematic representation of our workflow**, including the harmonisation, normalisation and integration of multiple genetic PD datasets, the generation of the IPD dataset, the development of the PD-KB based on data from experimental studies and public repositories, and the application of several statistical and computational methods (network-based analysis and pathway enrichment) towards IPD stratification. The colour of the arrows represents different methods applied in this study.

## Methods

Our workflow consisted of several complementary major steps within the systems biomedicine. This included harmonising and normalising multiple existing genetic PD datasets and integrating them into the PD-KG, along with data from public biological repositories. We also generated the IPD dataset and applied statistical and computation al methods for network-based analysis and pathway enrichment to help with IPD stratification. A summary of our methods is shown in Figure 6.

### Ethics approval

Written informed consent was obtained from all individuals who donated samples to this study. The work with iPSCs has been approved by the Ethics Review Panel (ERP) of the University of Luxembourg and the national Luxembourgish Research Ethics Committee (CNER, Comité National d’Ethique de Recherche) under the approval number CNER No. 201901/01 (ivPD).

### Midbrain organoid culture

Cell lines used in this study are summarised in Table M1. iPSCs were obtained from six healthy individuals (three female and three male individuals) and six age IPD patients (three female and three male individuals). Neuroepithelial stem cells (NESCs) derivation from iPSCs was performed as described in Reinhardt *et al.*, 2013^50^ via embryoid body formation and expansion of neuroectoderm. Midbrain organoids were generated and cultured as initially described in Monzel *et al.*, 2017^51^ (for image analysis) and further optimised by Nickels *et al.*, 2020^52^ (for RNA sequencing). In brief, NESCs at 80% of confluency were detached using Accutase (Sigma, Cat# A6964). Live cells were counted using Trypan Blue. 9x10 ^5^ cells were collected into 15ml of N2B27 maintenance media - DMEM-F12 (Thermo Fisher Scientific, cat.no 21331046) and Neurobasal (Thermo Fisher Scientific, Cat# 10888022) 50:50, supplemented with 1:200 N2 supplement (Thermo Fisher Scientific Cat# 17502001), 1:100 B27 supplement w/o Vitamin A (Life Technologies, Cat# 12587001), 1 % GlutaMAX (Thermo Fisher Scientific, Cat# 35050061) and 1 % penicillin/streptomycin (Thermo Fisher Scientific, Cat# 15140122). Cells were distributed in 96-well ultra-low attachment plates (faCellitate, Cat# F202003) - 150µl with 9000 cells per well. Plates were centrifuged at 300 g for one minute to facilitate spheroid formation at the bottom of the well. On the 2 ^nd^ day of organoid culture, media was changed to N2B27 pattering media, which is N2B27 base media supplemented with 10 ng/ml hBDNF (Peprotech, Cat# 450-02-1mg), 10 ng/ml hGDNF (Peprotech, Cat# 450-10-1mg), 500 µM dbcAMP (STEMCELL Technologies, Cat# 100-0244), 200 µM ascorbic acid (Sigma), 1 ng/ml TGF-β3 (Peprotech Cat# 100-36E) and 1 µM purmorphamine (Enzo Life Science, Cat# ALX-420-045). The next media change was done on the 5^th^ day of organoid culture. On day 8 of organoid culture, organoids for high-content imaging were embedded in extracellular matrix-like Geltrex (Thermo Fisher Scientific, Cat# A1413302) droplets^53^. Embedded organoids were kept in dynamic conditions on an orbital shaker (IKA), rotating at 80 rpm until the collection day. From day 8 of organoid culture organoids were kept in N2B27 differentiation media which only differs from the pattering media by lacking purmorphamine. Media changes were done every 3-4 days for embedded and non-embedded organoids until the day of sample collection. For single-cell sequencing, organoids were collected at day 50. For imaging, organoids were collected at days 30, 60, 90 and 180 of organoid culture. Cell culture was regularly tested for mycoplasma contamination using LookOut® Mycoplasma PCR Detection Kit (Sigma, Cat# MP0035-1KT).

### Immunofluorescence staining of organoid sections

Midbrain organoids were collected in a new 24-well plate (one organoid per well) and fixed with 4% paraformaldehyde (PFA) for six to seven hours at room temperature (RT) followed by washing three times with PBS for 15 minutes. Individual midbrain organoids were then embedded in 3% low-melting point agarose (Biozym Scientific GmbH, Cat# 840100). Midbrain organoids were sliced into 80 µm sections using a vibrating blade microtome (Leica VT1000s). Selected sections (per organoid one central section closer to the organoid core and one border section closer to the edge) for immunostaining were incubated for 30 min in 0.5% Triton X-100 at RT on an orbital shaker to permeabilize the cell membrane. Sections then were blocked for two hours at RT on an orbital shaker with a blocking buffer (2.5% normal donkey serum, 2.5 % BSA, 0.01% Triton X-100 and 0.1 % sodium azide). Primary antibodies were diluted in a blocking buffer and incubated with sections for 48 hours at 4 °C on an orbital shaker. Primary antibodies were stem cell markers PAX6 1:300 (Biolegend #901302, RRID: AB_2749901), SOX2 1:200 (R&D Systems Cat#BAF2018, RRID: AB_356217), KI67 1:200 (BD Biosciences Cat#550609, RRID: AB_393778), neuronal markers MAP2 1:1000 (Abcam Cat#ab5392, RRID: AB_2138153), TH 1:1000 (Abcam Cat#ab112, RRID: AB_297840), TUJ1 1:1 000 (BioLegend Cat#802001, RRID: AB_2564645 or Sigma-Aldrich Cat#AB9354, RRID:AB_570918), astrocyte markers S100beta 1:1000 (Sigma-Aldrich Cat#S2532, RRID: AB_477499), GFAP 1:1000 (Millipore Cat# AB5541, RRID:AB_177521) and alpha-Synclein 1:1000 (Novus Cat#NBP1-05194, RRID:AB_1555287). Then sections were washed three times for 5 min with 0.01% Triton X-100 in PBS followed by a 2-hour incubation at RT on an orbital shaker, protected from light with the Alexa Fluor^®^ conjugated secondary antibodies and nuclei stain Hoechst 33342 (Invitrogen Cat# 62249) diluted 1:1000 and 1:10 000 respectively in the blocking buffer. Sections were then washed three times for 10 min with 0.01% Triton X-100 in PBS and once with MiliQ water. For imaging sections were mounted on slides (De Beer Medicals, Cat# BM-9244) and covered with mounting media Fluoromount-G® (SouthernBiotech, Cat# 0100-01) and coverslip (VWR, Cat# ECN631-1574)

### Image acquisition and analysis

High-content imaging was performed using the Operetta high-content screening microscope (PerkinElmer) with a 20x objective using Z-stack acquisition selecting 25 planes per section. Acquired images were analysed with a customised pipeline using Matlab (v.2 017a, Mathworks, RRID: SCR_001622) as described in Bolognin *et al.*, 2018^54^ and Monzel *et al.*, 2020^55^. Only normalised features to the total nuclei count of the analysed organoids were considered in this study.

### Single-cell RNA sequencing and data analysis of IPD dataset

#### Midbrain organoids dissociation and single-cell isolation

Midbrain organoids were collected from their culture medium and washed with 1x PBS (phosphate-buffered saline, Gibco, Cat#10010-015). Organoids were transferred to a 1.5ml Eppendorf tube and digested in 1 ml sCelLive™ Tissue Dissociation Solution (Singleron Biotechnologies, Cat#1190062) diluted 1:2 with PBS. The organoids were placed in a thermal shaker at 750 rpm at 37°C for 45 minutes. The state of dissociation was checked at regular intervals under a light microscope. Following digestion, the suspension was filtered using a 40-µm sterile strainer (Greiner, Cat#542040). The cells were centrifuged at 350xg for 5 minutes at 4°C, and the cell pellets were resuspended in 300µl PBS. Cells were stained with Acridine Orange/Propidium Iodide Stain (Logos Biosystems, Cat#F23001), and the cell number and viability were calculated using LUNA-FX7™ Automated Cell Counter (Logos Biosystems, Villeneuve d’Ascq, France).

#### Single-cell RNA sequencing library Preparation

The single-cell RNA-seq libraries were constructed using GEXSCOPE™ Single Cell RNAseq Library Kit (Singleron Biotechnologies, Cat#4180011) according to the manufactureŕs instructions.

Briefly, for each library, the concentration of the single-cell suspension was adjusted to 3x105 cells/ml with PBS, and the suspension was loaded onto an SD microfluidic chip to capture 6000 cells. Paramagnetic beads conjugated to oligodT probes that carry a unique molecular identifier (UMI) and a barcode unique to each bead (from the same kit) were loaded, after which the cells were lysed. The beads bound to polyadenylated mRNA were extracted from the chip and reverse transcribed into cDNA at 42°C for 1.5 hours, and the cDNA was amplified by PCR. The cDNA was then fragmented and ligated to indexed Illumina adapters. The fragment size distribution of the final amplified library was obtained on an Agilent TapeStation.

#### Library sequencing

The library concentration was calculated using the Qubit 4.0 fluorometer and the libraries were pooled in an equimolar fashion. The single cell libraries were sequenced on an Illumina NovaSeq X using a 2x150-bp approach to a final depth of 90 GB per library. The reads were demultiplexed according to the multiplexing index sequencing on Illumina’s BaseCloud platform.

#### Transcriptome data pre-processing

The pre-processing of the fastq files was conducted using CeleScopeÂ® (v.1.14.1; www.github.com/singleron-RD/CeleScope; Singleron Biotechnologies GmbH) to generate gene count matrices with default parameters. Low-quality reads were removed. Sequences were mapped using STAR (https://github.com/alexdobin/STAR) to the human genome reference version GRCh38 and genes were annotated using Ensembl 92. The reads were assigned to genes using featureCount (https://subread.sourceforge.net/) and the cell calling was performed by fitting a negative bimodal distribution and determining the threshold between empty wells and cell-associated wells to generate a count matrix file containing the number of Unique Molecular Identifier (UMI) for each gene within each cell.

#### Transcriptome data analysis

Downstream analysis was done using Seurat single-cell analysis toolkit (v. 5.0.1; RRID: SCR_016341)^56^ in R computing language (v. 4.3.2; RRID: SCR_001905). Cells with less than 500 genes and more than 6000 -7500 (customised setting depending on the sample) were excluded to remove non-viable or low-quality cells and doublets respectively. In addition, cells having a mitochondrial gene content above 15-20% were also excluded as non-viable cells. Datasets of single cell lines were merged, log normalised and scaled to all genes. The integration was performed using Seurat integration workflow 5.0 ^56^ based on the first 20 PCA components. Cell populations were identified by applying the Louvain algorithm modularity optimization with a resolution of 0.2^57^. Nine distinct cell clusters were identified and visualised using the uniform manifold approximation and projection (UMAP) technique ^58^. Marker genes of each cell population were determined by applying the *FindAllMarkers* function of Seurat. Additionally, cellular identities were validated using the GeneAnalytics online tool ^59^ (https://geneanalytics.genecards.org/), PangloaDB^36^ (https://panglaodb.se/) and human midbrain cell type-specific markers reported in La Manno *et al.*, 2016^35^.

### Data integration and *in silico* analysis

#### PD data corpus creation from existing PD datasets

First, we created a core data corpus considering the top 100 most significant DEGs (selected by the adjusted p.value from available bulk or single cell-RNA sequencing data) and 12 core proteins selected from high-content imaging data of four independent published datasets of midbrain organoids (Table M2). Long-non-coding RNAs and pseudogenes were excluded. The original RNA sequencing experiments were bulk for the GBA-PD and SNCA-PD and single-cell for MIRO1-PD and LRRK2-PD mutations. In the latter case, the top 100 significant

**Table M2.** Previously published studies using midbrain organoids integrated to create the graph database.

| Mutation | Study | Main findings |
| --- | --- | --- |
| **LRRK2-G2019S** | Smits *et al.*, 2019 (imaging)<sup>28</sup><br>Smits *et al.*, 2020 (RNA seq)<sup>29</sup> | - Neurodevelopmental defects of midbrain dopaminergic neurons |
| **3xSNCA** | Modamio *et al.*, 2021<sup>30</sup> | - Increased levels of α-synuclein and, phosphorylated (pSer129) α-synuclein<br>- Impaired synaptic function<br>- Loss of dopaminergic neurons over time |
| **GBA-N370S** | Rosety *et al.*, 2023<sup>31</sup> | - Neurogenesis defects<br>- Altered lipid metabolism,<br>- Reduced GCase activity<br>- Reduced levels of dopaminergic neurons |
| **MIRO1-R272Q** | Chemla *et al.*, 2023<sup>32</sup> | - Mitochondrial impairments<br>- Increased levels of α-synuclein<br>- Dopaminergic neuron death |

DEGs were determined in a bulk mode to enable comparison to the GBA-PD and SNCA-PD RNA sequencing results. For PCA and unsupervised hierarchical clustering on different PD datasets log2 fold changes of PD vs control samples from every individual RNA sequencing experiment were considered. For PCA on IPD vs control samples from the same dataset, transcriptomics data was integrated as the average gene expression across all cells. Stratification analysis was performed in R (v. 4.3.2; RRID: SCR_001905). Imaging datasets from four previously published studies were harmonised defining common column names to enable downstream analysis.

#### PD-KG development

We developed the PD-KG by using the neo4j technology (neo4j-community-4.4.25 version) and the Java Eclipse environment (with Java SE 1.8). At the core, the PD-KB integrates molecular features from high-content imaging and transcriptomics data from PD iPSCs-generated midbrain organoids. We focused on the contextualisation of these molecular features from the experimental datasets. We integrated information on several types of biological relationships involving these molecular features and other biological entities from public biological repositories such as i) pathway involvement from Reactome (filtered on the human species)^23^, ii) disease association from DisGeNet^25^, iii) drug targets from DGIdb^26^, iv) protein-protein association from IntAct (we selected several types of interactions, including association, physical association, colocalization and proximity) ^24^ . The mapping between the HGNC gene symbols and unique UNIPROT identifiers was done using UniProtKB^27^. The reference date for data integration was 04/06/2024. In the underlying graph, the data types such as core proteins, genes, disease, drugs, and pathways were represented by nodes, and their inter-relationships were shown as connecting edges. The graph data model is shown in Figure 1 and the types of data and their inter-relationships are given in Supplementary Data.

#### Network-based analysis

We performed network-based analysis on the PD data corpus at several levels as follows:

#### Similarity Network Fusion (SNF) for the idiopathic PD dataset

We performed a SNF analysis^38^ towards the IPD cell line stratification by integrating imaging data and top 100 transcriptomics from 6 IPD cell lines and 3 healthy controls, respectively. The transcriptomics data was integrated as the average gene expression across all cells. The code was developed in RStudio (version 2024.04.2+764) under the following settings: R version 4.1.0 (2021-05-18), Platform: x86_64-apple-darwin17.0 (64-bit), running under: macOS 14.5. We used the SNFtool package for the SNF and the spectralClustering methods, the pheatmap package for the heatmap of the results.

#### Over-representation analysis (ORA) for the genetic and idiopathic PD datasets

We performed an over-representation analysis (ORA)^33^ in R version 4.1.0 (2021-05-18) for the top 100 transcriptomics features selected from the existing genetic PD and IPD datasets, respectively. We checked the enriched sets of Reactome pathways corresponding to each of the input gene datasets. We used the following methods and packages in RStudio, (with the same settings as for the SNF analysis):

- enrichPathway and ReactomePA for the Reactome pathway enrichment;
- org.Hs.eg.db and AnnotationDbi::select for the gene ID conversion (between the gene symbol and Entrez id);
- dotplot for the visualisation of the enrichment results;
- ggtitle, theme and ggplot2 for the final customised plots of the enrichment results.

#### Core analysis using gene expression for pathway activity prediction

We computed the set of top 100 DEG for every IPD cell line in comparison with the healthy samples and performed a core analysis in the Ingenuity Pathway Analysis platform (IPA)^39^ to predict the list of canonical pathways being activated or inhibited in the IPD samples. We used IPA Version 01-21-02, with the following settings for the core analysis: Species = human, Reference set = “Ingenuity Knowledge Base (Gene Only”), Relationships to consider = “Direct Relationships”.

## Data availability

All data supporting the conclusions of this manuscript are publicly available under this DOI: 10.17881/xs23-rk90.

RNA sequencing datasets are available on Gene Expression Omnibus (GEO) under the accession codes: GSE237133 (MIRO1-PD), GSE133894 (LRRK2-PD), GSE208784 (GBA-PD), GSE278265 (SNCA-PD), GSE276684 (IPD).

## Code availability

All computational scripts used for the modelling and analysis are available on GitLab at: https://gitlab.com/uniluxembourg/lcsb/developmental-and-cellular-biology/zagare_tandem_2024

## Acknowledgements

We acknowledge funding from LCSB Tandem Project 2023 for the IPD single -cell RNA sequencing dataset generation. We thank the Pre-Publication Check (PPC) team at LCSB-UNILU for the comments and assistance with ensuring the FAIRness and reproducibility of this work. We would also like to acknowledge access to the Neo4j framework and to public repositories: DisGeNET, DGIdb, IntAct, Reactome, UniProt. We thank Dr. Nico J. Diederich and Laura Longhino from Centre Hospitalier de Luxembourg, Thomas Rauen, Sergii Velychko, Anna-Lena Hallmann and Hans Schoeler from the Max Planck Institute in Muenster, Dr. Kathleen Mommaerts from Integrated BioBank of Luxembourg, Dr. Bill Skarnes and the Coriell Institute for providing cell lines. Further, we acknowledge Dr. Isabel Rosety, Dr. Lisa Smits and Dr. Jennifer Modamio for the generation of the high-content imaging and RNA sequencing data. High-content imaging and customised script optimisation were supported by the LCSB bio-imaging platform and Dr. Paul Antony. Single-cell RNA sequencing was done by Singleron Biotechnologies.

## Author contributions

Conceptualisation: AZ, IB, CS, MG, SG, VPS, JCS. Methodology: AZ, IB, CS, MG, SG, AR ASM. Formal analysis: AZ, IB, AR, MG, SG. Computational analysis/scripts: IB, AZ, AR, SG. Initial draft preparation: AZ, IB. Review and editing: all authors. Project coordination: VPS and JCS. All authors have read and agreed to the final version of the manuscript.

## Rights retention statement

“For the purpose of Open Access, the author has applied a CC BY public copyright licence to any Author Accepted Manuscript (AAM) version arising from this submission.”

