## Supplementary Figures for "Deciphering shared molecular dysregulation across Parkinson’s Disease variants using a multi-modal network-based data integration and analysis"

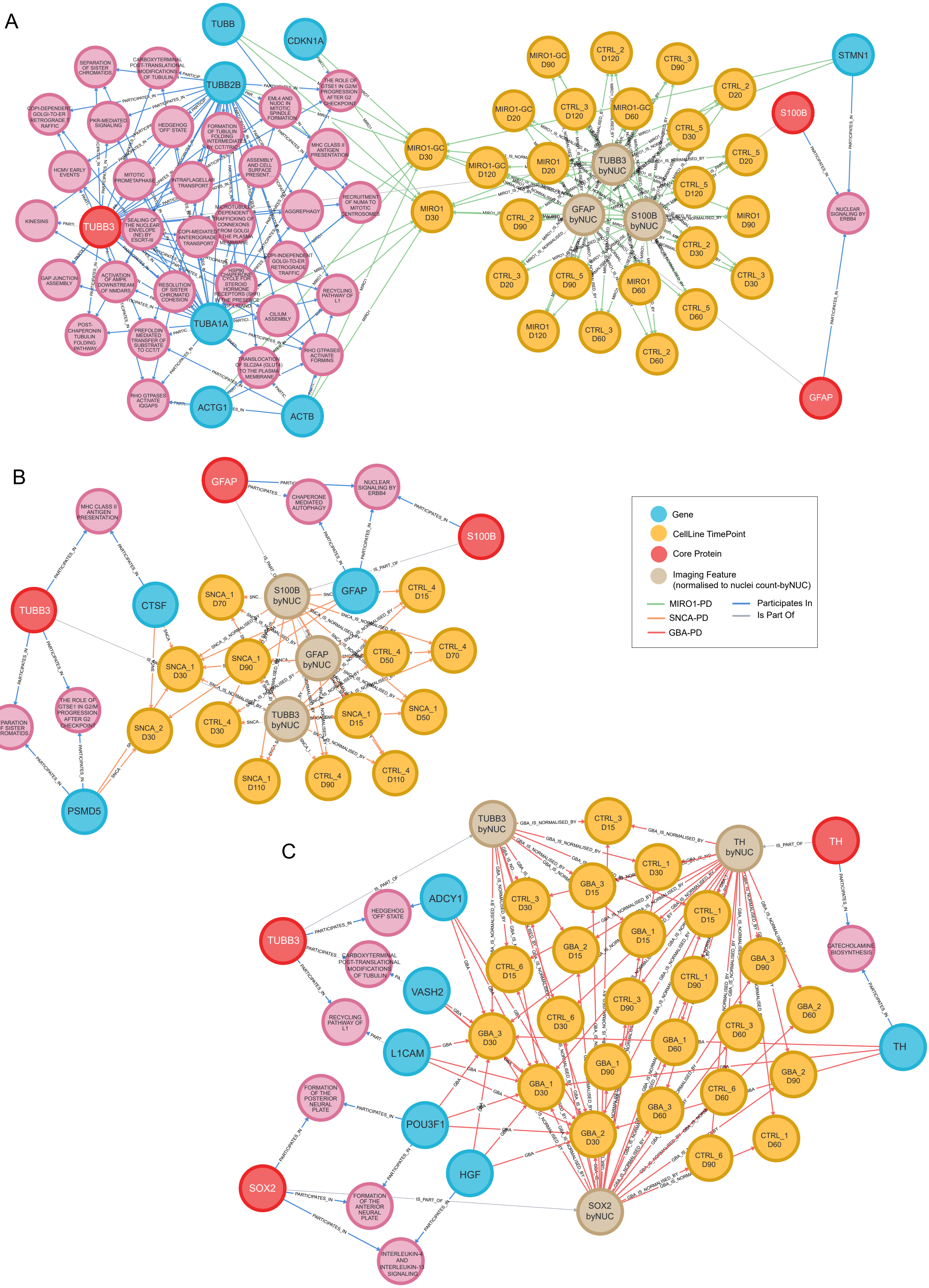


**Figure S1.** Relationships between core proteins and the top significantly differentially expressed genes in individual datasets A) MIRO1-PD, B) SNCA-PD and C) GBA-PD.

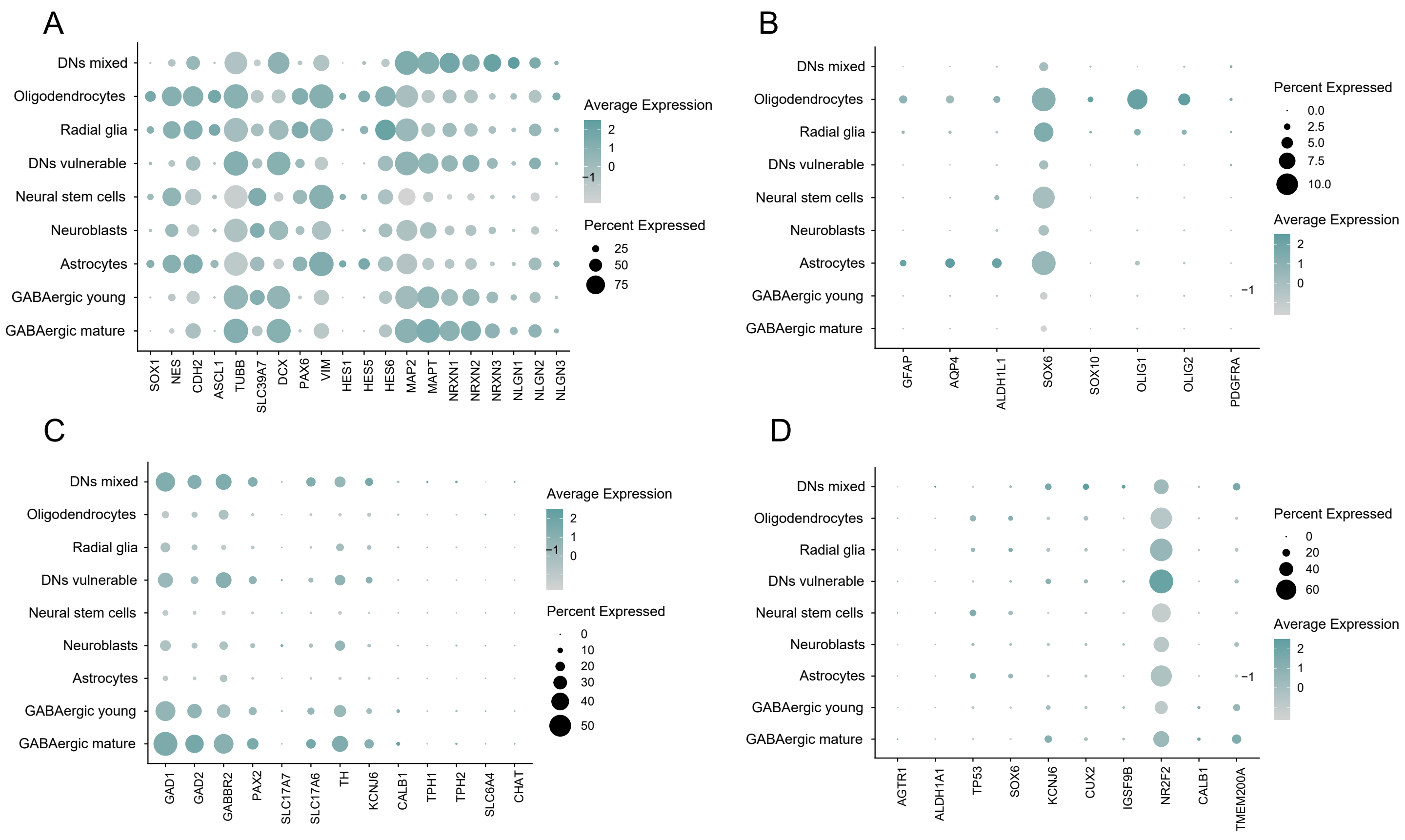


**Figure S2.** Cell type marker expression. A) Neuronal stem cell (SOX1, NES), neuroblast (CDH2, ASCL1, TUBB,SLC39A7, DCX, PAX6), radial glia (VIM, HES1, HES5, HES6) and mature neuron markers (MAP2, MAPT, NRXN1, NRXN2, NRXN3, NLGN1, NLGN2, NLGN3). B) Atrocyte (GFAP, AQP4, ALDH1L1, SOX6) and oligodendrocyte (SOX6, SOX10, OLIG1, OLIG2) markers. Oligodendrocyte progenitor cell marker PDGFRA to confirm oligodendrocyte maturity. C) Neuron subtype markers: GABAergic neurons (GAD1, GAD2, GABBRA2), glutamatergic neurons (PAX2, SLC17A7, SLC17A6), dopaminergic neurons (TH, KCNJ6, CALB1), serotonergic neurons (TPH1, TPH2, SLC6A4) and cholinergic neurons (CHAT). D) Dopaminergic neurons vulnerability (AGTR1, ALDH1A1, TP53, SOX6, KCNJ6, CUX2, IGSF9B, NRF2F2) and resistance (CALB1, TMEM200A) markers.

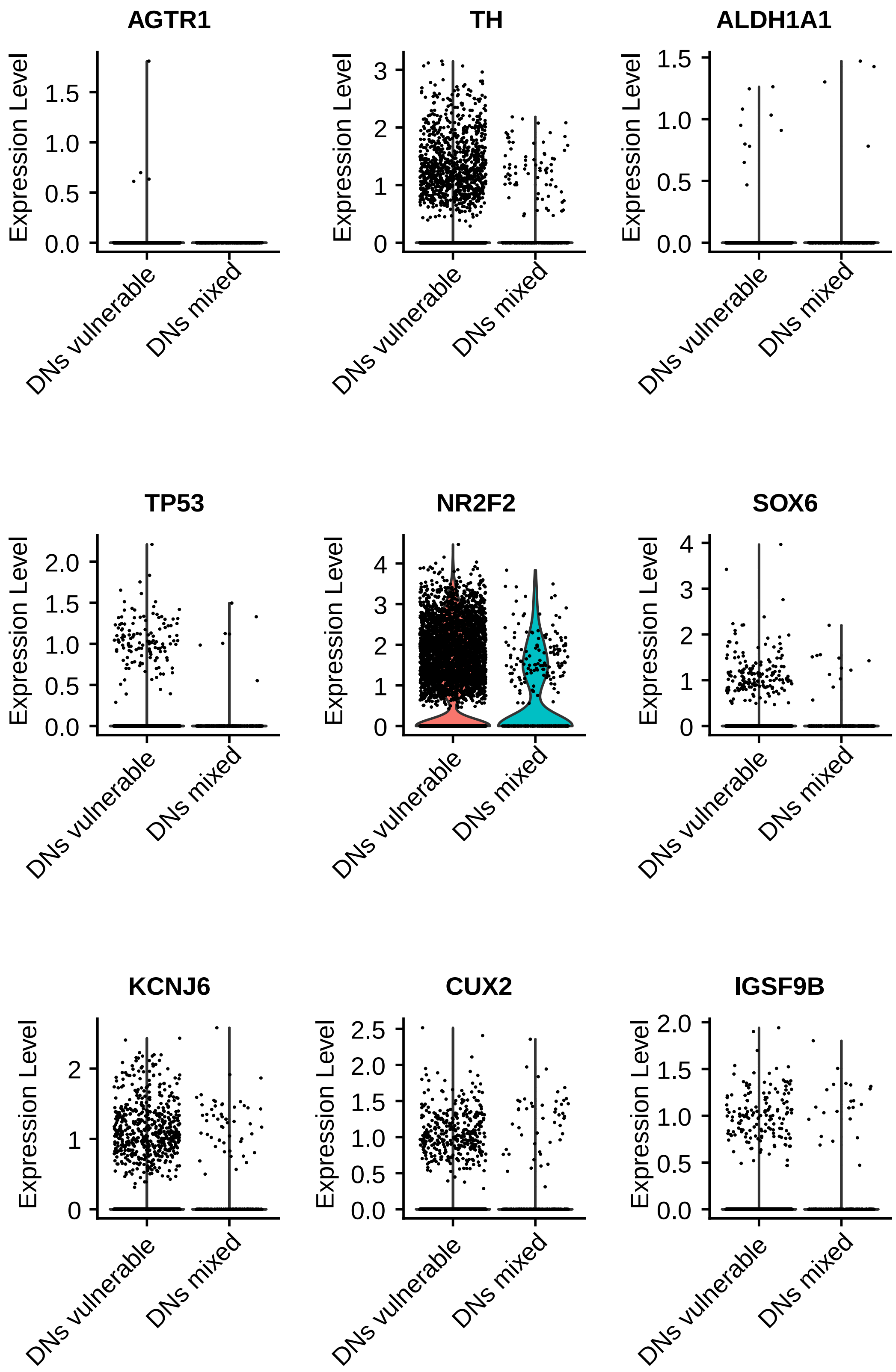


**Figure S3.** Expression of dopaminergic neuron vulnerability markers in the two dopaminergic neuron populations.

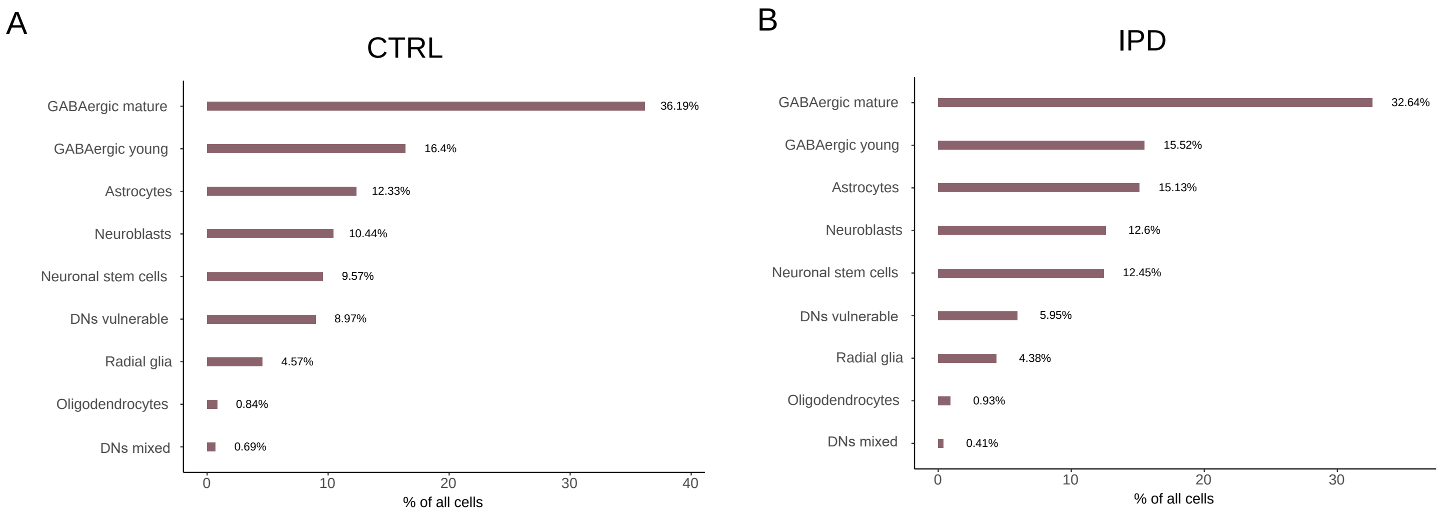


**Figure S4.** Percentage of all cellular populations calculated from the total amount of cells in CTRL and IPD samples.

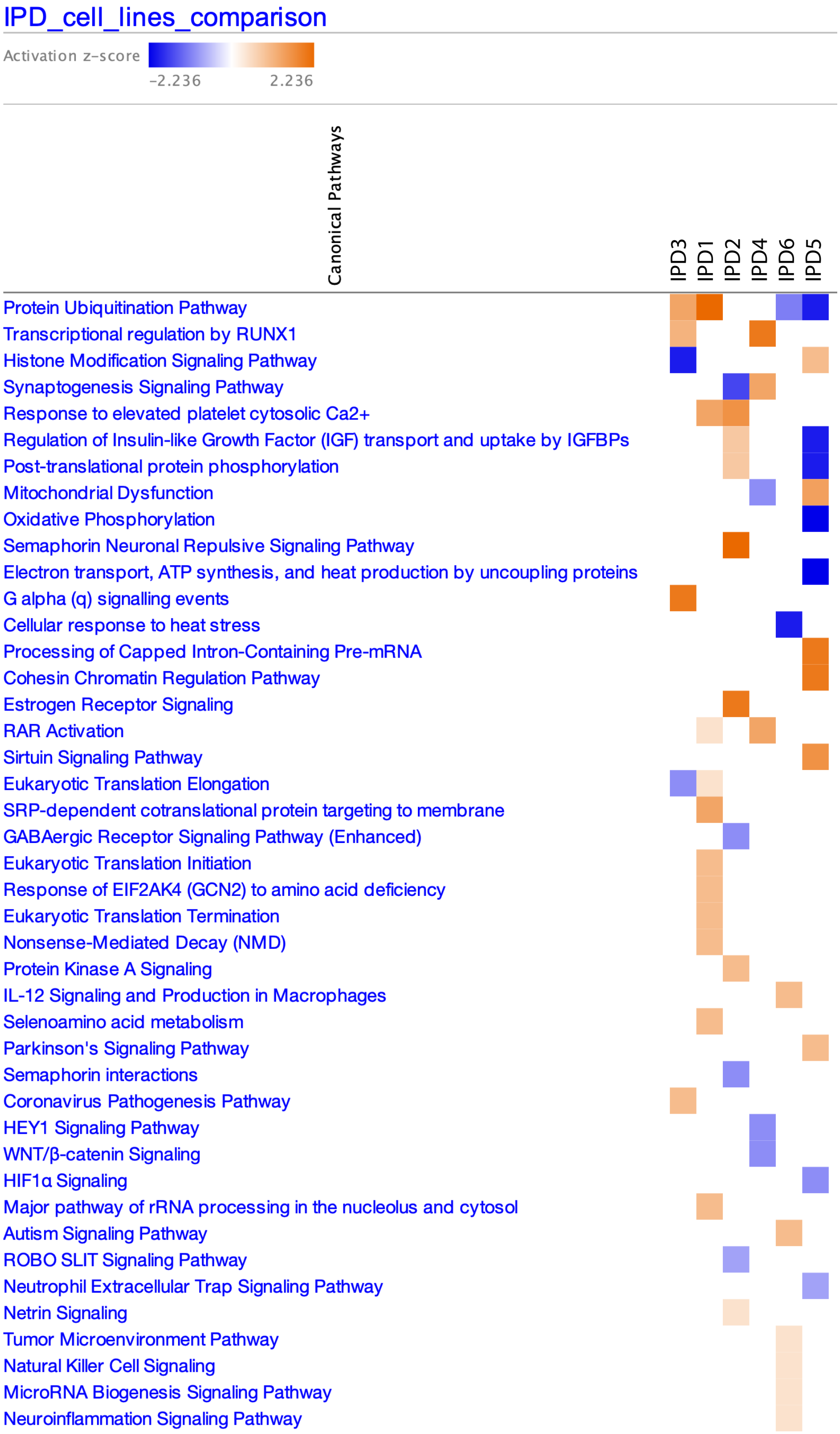


**Figure S5.** Ingenuity pathway analysis to predict the activation level of metabolic and signaling pathways based on log2 fold change (IPD vs CTRL) of the top 100 significant differential expressed genes determined for each IPD lines against all six CTRL samples.
