## Supplementary Tables for "Deciphering shared molecular dysregulation across Parkinson’s Disease variants using a multi-modal network-based data integration and analysis"

**Table S1.** List of 25 genes shared between at least two RNA sequencing experiments of monogenic PD patient specific midbrain organoid studies

| Gene symbol | No. of shared experiments | Shared experiments |
|---|---|---|
| FTL | 2 | LRRK2, MIRO1 |
| PTPRD | 2 | LRRK2, MIRO1 |
| MAB21L2 | 2 | LRRK2, GBA |
| GAD1 | 2 | LRRK2, MIRO1 |
| GRIA2 | 2 | LRRK2, GBA |
| CHL1 | 2 | LRRK2, MIRO1 |
| TUBA1A | 2 | LRRK2, MIRO1 |
| ACTB | 2 | LRRK2, MIRO1 |
| ACTG1 | 2 | LRRK2, MIRO1 |
| ENO1 | 2 | LRRK2, MIRO1 |
| PAX5 | 2 | LRRK2, SNCA |
| MORF4L2 | 2 | LRRK2, MIRO1 |
| MAB21L1 | 2 | LRRK2, GBA |
| IGFBP2 | 2 | LRRK2, MIRO1 |
| EN1 | 2 | LRRK2, MIRO1 |
| SLC17A6 | 2 | LRRK2, MIRO1 |
| TUBB2B | 2 | LRRK2, MIRO1 |
| OTP | 2 | LRRK2, SNCA |
| ROBO2 | 2 | LRRK2, MIRO1 |
| PEG10 | 2 | LRRK2, MIRO1 |
| EPHA5 | 2 | GBA, MIRO1 |
| TCEAL7 | 2 | GBA, MIRO1 |
| TMEM132B | 2 | GBA, SNCA |
| CBLN1 | 2 | SNCA, MIRO1 |
| DPP6 | 2 | SNCA, MIRO1 |

**Table S2.** Expression pattern of the 25 shared genes between at least two experiments in PD samples vs healthy controls (CTRL).

| Gene symbol | Expression in PD vs CTRL | Shared experiments |
|---|---|---|
| MAB21L2 | DOWN | LRRK2, GBA |
| GAD1 | DOWN | LRRK2, MIRO1 |
| GRIA2 | DOWN | LRRK2, GBA |
| TUBA1A | UP | LRRK2, MIRO1 |
| ACTB | UP | LRRK2, MIRO1 |
| ACTG1 | UP | LRRK2, MIRO1 |
| MAB21L1 | DOWN | LRRK2, GBA |
| TUBB2B | UP | LRRK2, MIRO1 |
| PEG10 | UP | LRRK2, MIRO1 |
| TCEAL7 | DOWN | GBA, MIRO1 |
| CBLN1 | UP | SNCA, MIRO1 |
| DPP6 | UP | SNCA, MIRO1 |
| FTL | UP, DOWN | LRRK2, MIRO1 |
| PTPRD | DOWN, UP | LRRK2, MIRO1 |
| CHL1 | DOWN, UP | LRRK2, MIRO1 |
| ENO1 | UP, DOWN | LRRK2, MIRO1 |
| PAX5 | DOWN, UP | LRRK2, SNCA |
| MORF4L2 | UP, DOWN | LRRK2, MIRO1 |
| IGFBP2 | UP, DOWN | LRRK2, MIRO1 |
| EN1 | UP, DOWN | LRRK2, MIRO1 |
| SLC17A6 | DOWN, UP | LRRK2, MIRO1 |
| OTP | DOWN, UP | LRRK2, SNCA |
| ROBO2 | DOWN, UP | LRRK2, MIRO1 |
| EPHA5 | DOWN, UP | GBA, MIRO1 |
| TMEM132B | DOWN, UP | GBA, SNCA |

**Table S3.** Reactome pathways associated with the most connected six out of 25 genes shared between genetic PD datasets.

| Pathway | Gene symbol | No. of genes |
|---|---|---|
| "RECYCLING PATHWAY OF L1" | ["ACTB", "TUBA1A", "ACTG1", "TUBB2B"] | 4 |
| "TRANSLOCATION OF SLC2A4 (GLUT4) TO THE PLASMA MEMBRANE" | ["ACTG1", "TUBB2B", "ACTB", "TUBA1A"] | 4 |
| "RHO GTPASES ACTIVATE IQGAPS" | ["ACTG1", "TUBB2B", "TUBA1A", "ACTB"] | 4 |
| "RHO GTPASES ACTIVATE FORMINS" | ["ACTB", "TUBB2B", "TUBA1A", "ACTG1"] | 4 |
| "EPH-EPHRIN MEDIATED REPULSION OF CELLS" | ["EPHA5", "ACTB", "ACTG1"] | 3 |
| "PREFOLDIN MEDIATED TRANSFER OF SUBSTRATE TO CCT/TRIC" | ["TUBB2B", "TUBA1A", "ACTB"] | 3 |
| "RECRUITMENT OF NUMA TO MITOTIC CENTROSOMES" | ["TUBB2B", "TUBA1A"] | 2 |
| "VEGFA-VEGFR2 PATHWAY" | ["ACTG1", "ACTB"] | 2 |
| "CELL-EXTRACELLULAR MATRIX INTERACTIONS" | ["ACTB", "ACTG1"] | 2 |
| "ASSEMBLY AND CELL SURFACE PRESENTATION OF NMDA RECEPTORS" | ["TUBA1A", "TUBB2B"] | 2 |
| "HCMV EARLY EVENTS" | ["TUBA1A", "TUBB2B"] | 2 |
| "INTERACTION BETWEEN L1 AND ANKYRINS" | ["ACTG1", "ACTB"] | 2 |
| "ACTIVATION OF AMPK DOWNSTREAM OF NMDARS" | ["TUBB2B", "TUBA1A"] | 2 |
| "SEPARATION OF SISTER CHROMATIDS" | ["TUBA1A", "TUBB2B"] | 2 |
| "SIGNALING DOWNSTREAM OF RAS MUTANTS" | ["ACTB", "ACTG1"] | 2 |
| "CLATHRIN-MEDIATED ENDOCYTOSIS" | ["ACTG1", "ACTB"] | 2 |
| "SIGNALING BY RAF1 MUTANTS" | ["ACTG1", "ACTB"] | 2 |
| "SEALING OF THE NUCLEAR ENVELOPE (NE) BY ESCRT-III" | ["TUBB2B", "TUBA1A"] | 2 |
| "ADHERENS JUNCTIONS INTERACTIONS" | ["ACTB", "ACTG1"] | 2 |
| "SIGNALING BY BRAF AND RAF1 FUSIONS" | ["ACTG1", "ACTB"] | 2 |
| "PARADOXICAL ACTIVATION OF RAF SIGNALING BY KINASE INACTIVE BRAF" | ["ACTG1", "ACTB"] | 2 |
| "THE ROLE OF GTSE1 IN G2/M PROGRESSION AFTER G2 CHECKPOINT" | ["TUBA1A", "TUBB2B"] | 2 |
| "CILIUM ASSEMBLY" | ["TUBA1A", "TUBB2B"] | 2 |
| "SIGNALING BY MODERATE KINASE ACTIVITY BRAF MUTANTS" | ["ACTB", "ACTG1"] | 2 |

| | | |
|---|---|---|
| **"SIGNALING BY HIGH-KINASE ACTIVITY BRAF MUTANTS"** | ["ACTG1", "ACTB"] | 2 |
| **"EPHB-MEDIATED FORWARD SIGNALING"** | ["ACTB", "ACTG1"] | 2 |
| **"REGULATION OF ACTIN DYNAMICS FOR PHAGOCYTIC CUP FORMATION"** | ["ACTG1", "ACTB"] | 2 |
| **"COPI-INDEPENDENT GOLGI-TO-ER RETROGRADE TRAFFIC"** | ["TUBA1A", "TUBB2B"] | 2 |
| **"COPI-DEPENDENT GOLGI-TO-ER RETROGRADE TRAFFIC"** | ["TUBA1A", "TUBB2B"] | 2 |
| **"KINESINS"** | ["TUBB2B", "TUBA1A"] | 2 |
| **"AGGREPHAGY"** | ["TUBB2B", "TUBA1A"] | 2 |
| **"MAP2K AND MAPK ACTIVATION"** | ["ACTG1", "ACTB"] | 2 |
| **"MHC CLASS II ANTIGEN PRESENTATION"** | ["TUBA1A", "TUBB2B"] | 2 |
| **"RESOLUTION OF SISTER CHROMATID COHESION"** | ["TUBB2B", "TUBA1A"] | 2 |
| **"MITOTIC PROMETAPHASE"** | ["TUBB2B", "TUBA1A"] | 2 |
| **"PKR-MEDIATED SIGNALING"** | ["TUBA1A", "TUBB2B"] | 2 |
| **"INTRAFLAGELLAR TRANSPORT"** | ["TUBA1A", "TUBB2B"] | 2 |
| **"SENSORY PROCESSING OF SOUND BY INNER HAIR CELLS OF THE COCHLEA"** | ["ACTB", "ACTG1"] | 2 |
| **"SENSORY PROCESSING OF SOUND BY OUTER HAIR CELLS OF THE COCHLEA"** | ["ACTG1", "ACTB"] | 2 |
| **"FCGR3A-MEDIATED PHAGOCYTOSIS"** | ["ACTG1", "ACTB"] | 2 |
| **"GAP JUNCTION DEGRADATION"** | ["ACTG1", "ACTB"] | 2 |
| **"HEDGEHOG 'OFF' STATE"** | ["TUBA1A", "TUBB2B"] | 2 |
| **"GAP JUNCTION ASSEMBLY"** | ["TUBB2B", "TUBA1A"] | 2 |
| **"FORMATION OF ANNULAR GAP JUNCTIONS"** | ["ACTB", "ACTG1"] | 2 |
| **"MICROTUBULE-DEPENDENT TRAFFICKING OF CONNEXONS FROM GOLGI TO THE PLASMA MEMBRANE"** | ["TUBB2B", "TUBA1A"] | 2 |
| **"CARBOXYTERMINAL POST-TRANSLATIONAL MODIFICATIONS OF TUBULIN"** | ["TUBA1A", "TUBB2B"] | 2 |
| **"POST-CHAPERONIN TUBULIN FOLDING PATHWAY"** | ["TUBB2B", "TUBA1A"] | 2 |
| **"HATS ACETYLATE HISTONES"** | ["ACTB", "MORF4L2"] | 2 |
| **"COPI-MEDIATED ANTEROGRADE TRANSPORT"** | ["TUBB2B", "TUBA1A"] | 2 |
| **"EML4 AND NUDC IN MITOTIC SPINDLE FORMATION"** | ["TUBA1A", "TUBB2B"] | 2 |

| | | |
|---|---|---|
| **"HSP90 CHAPERONE CYCLE FOR STEROID HORMONE RECEPTORS (SHR) IN THE PRESENCE OF LIGAND"** | ["TUBB2B", "TUBA1A"] | 2 |
| **"RHO GTPASES ACTIVATE WASPS AND WAVES"** | ["ACTB", "ACTG1"] | 2 |
| **"FORMATION OF TUBULIN FOLDING INTERMEDIATES BY CCT/TRIC"** | ["TUBA1A", "TUBB2B"] | 2 |
| **"RHOBTB2 GTPASE CYCLE"** | ["ACTG1"] | 1 |
| **"B-WICH COMPLEX POSITIVELY REGULATES RRNA EXPRESSION"** | ["ACTB"] | 1 |
| **"REGULATION OF MITF-M-DEPENDENT GENES INVOLVED IN PIGMENTATION"** | ["ACTB"] | 1 |
| **"FOLDING OF ACTIN BY CCT/TRIC"** | ["ACTB"] | 1 |
| **"FACTORS INVOLVED IN MEGAKARYOCYTE DEVELOPMENT AND PLATELET PRODUCTION"** | ["ACTB"] | 1 |
| **"ANCHORING OF THE BASAL BODY TO THE PLASMA MEMBRANE"** | ["TUBA1A"] | 1 |
| **"LOSS OF PROTEINS REQUIRED FOR INTERPHASE MICROTUBULE ORGANIZATION FROM THE CENTROSOME"** | ["TUBA1A"] | 1 |
| **"EPH-EPHRIN SIGNALING"** | ["EPHA5"] | 1 |
| **"RHOF GTPASE CYCLE"** | ["ACTB"] | 1 |
| **"RECRUITMENT OF MITOTIC CENTROSOME PROTEINS AND COMPLEXES"** | ["TUBA1A"] | 1 |
| **"LOSS OF NLP FROM MITOTIC CENTROSOMES"** | ["TUBA1A"] | 1 |
| **"EPHA-MEDIATED GROWTH CONE COLLAPSE"** | ["EPHA5"] | 1 |
| **"DNA DAMAGE RECOGNITION IN GG-NER"** | ["ACTB"] | 1 |
| **"AURKA ACTIVATION BY TPX2"** | ["TUBA1A"] | 1 |
| **"REGULATION OF PLK1 ACTIVITY AT G2/M TRANSITION"** | ["TUBA1A"] | 1 |
| **"UCH PROTEINASES"** | ["ACTB"] | 1 |

**Table S4.** List of Core Proteins. Proteins that were immunostained in different experiments for high-content imaging analysis.

| No. | Gene symbol | Uniprot Id | Protein name | Experiments |
|---|---|---|---|---|
| **1** | CNP | P09543 | 2',3'-cyclic-nucleotide 3'-phosphodiesterase | GBA |
| **2** | DCX | O43602 | Neuronal migration protein doublecortin | GBA |
| **3** | FOXA2 | Q9Y261 | Hepatocyte nuclear factor 3-beta | GBA, LRRK2 |
| **4** | GFAP | P14136 | Glial fibrillary acidic protein | SNCA, MIRO1, IPD, GBA |
| **5** | MAP2 | P11137 | Microtubule-associated protein 2 | SNCA, MIRO1, IPD, GBA |
| **6** | MKI67 | P46013 | Proliferation marker protein Ki-67 | IPD, GBA |
| **7** | NES | P48681 | Nestin | GBA |
| **8** | OLIG2 | Q13516 | Oligodendrocyte transcription factor 2 | GBA |
| **9** | PAX6 | P26367 | Paired box protein Pax-6 | IPD |
| **10** | S100B | P04271 | Protein S100-B | SNCA, MIRO1, IPD, GBA |
| **11** | SOX2 | P48431 | Transcription factor SOX-2 | GBA, IPD |
| **12** | TH | P07101 | Tyrosine hydroxylase | SNCA, MIRO1, IPD, GBA, LRRK2 |
| **13** | TUBB3/TUJ1 | Q13509 | Tubulin beta-3 chain | [SNCA, MIRO1, IPD, GBA] |

**Table S5.** Shared pathways between core proteins and significant differential expressed genes in individual datasets

| Reactome Pathway name | Core proteins | Genes | Experiment |
|---|---|---|---|
| CARBOXYTERMINAL POST-TRANSLATIONAL MODIFICATIONS OF TUBULIN | TUBB3 | VASH2 | GBA |
| CARBOXYTERMINAL POST-TRANSLATIONAL MODIFICATIONS OF TUBULIN | TUBB3 | TUBA1A, TUBB2B | MIRO1 |
| HEDGEHOG 'OFF' STATE | TUBB3 | ADCY1 | GBA |
| HEDGEHOG 'OFF' STATE | TUBB3 | TUBA1A, TUBB2B | MIRO1 |
| RECYCLING PATHWAY OF L1 | TUBB3 | L1CAM | GBA |
| RECYCLING PATHWAY OF L1 | TUBB3 | ACTB, TUBA1A, ACTG1, TUBB2B | MIRO1 |
| SEPARATION OF SISTER CHROMATIDS | TUBB3 | PSMD5 | SNCA |
| SEPARATION OF SISTER CHROMATIDS | TUBB3 | TUBA1A, TUBB2B | MIRO1 |
| THE ROLE OF GTSE1 IN G2/M PROGRESSION AFTER G2 CHECKPOINT | TUBB3 | PSMD5 | SNCA |
| THE ROLE OF GTSE1 IN G2/M PROGRESSION AFTER G2 CHECKPOINT | TUBB3 | TUBA1A, CDKN1A, TUBB2B | MIRO1 |
| MHC CLASS II ANTIGEN PRESENTATION | TUBB3 | CTSF | SNCA |
| MHC CLASS II ANTIGEN PRESENTATION | TUBB3 | TUBA1A, TUBB2B | MIRO1 |
| NUCLEAR SIGNALING BY ERBB4 | GFAP, S100B | GFAP | SNCA |
| NUCLEAR SIGNALING BY ERBB4 | GFAP, S100B | STMN1 | MIRO1 |
| CATECHOLAMINE BIOSYNTHESIS | TH | TH | GBA |
| CHAPERONE MEDIATED AUTOPHAGY | GFAP | GFAP | SNCA |
| FORMATION OF THE ANTERIOR NEURAL PLATE | SOX2 | POU3F1 | GBA |
| FORMATION OF THE POSTERIOR NEURAL PLATE | SOX2 | POU3F1 | GBA |
| INTERLEUKIN-4 AND INTERLEUKIN-13 SIGNALING | SOX2 | HGF | GBA |
| COPI-DEPENDENT GOLGI-TO-ER RETROGRADE TRAFFIC | TUBB3 | TUBA1A, TUBB2B | MIRO1 |
| RECRUITMENT OF NUMA TO MITOTIC CENTROSOMES | TUBB3 | TUBB2B, TUBA1A, TUBB | MIRO1 |
| CILIUM ASSEMBLY | TUBB3 | TUBA1A, TUBB2B | MIRO1 |
| INTRAFLAGELLAR TRANSPORT | TUBB3 | TUBA1A, TUBB2B | MIRO1 |
| TRANSLOCATION OF SLC2A4 (GLUT4) TO THE PLASMA MEMBRANE | TUBB3 | ACTG1, TUBB2B, ACTB, TUBA1A | MIRO1 |
| HSP90 CHAPERONE CYCLE FOR STEROID HORMONE RECEPTORS (SHR) IN THE PRESENCE OF LIGAND | TUBB3 | TUBB2B, TUBA1A | MIRO1 |
| RHO GTPASES ACTIVATE FORMINS | TUBB3 | ACTB, TUBB2B, TUBA1A, ACTG1 | MIRO1 |
| ACTIVATION OF AMPK DOWNSTREAM OF NMDARS | TUBB3 | TUBB2B, TUBA1A | MIRO1 |
| SEALING OF THE NUCLEAR ENVELOPE (NE) BY ESCRT-III | TUBB3 | TUBB2B, TUBA1A | MIRO1 |
| RHO GTPASES ACTIVATE IQGAPS | TUBB3 | ACTG1, TUBB2B, TUBA1A, ACTB | MIRO1 |
| FORMATION OF TUBULIN FOLDING INTERMEDIATES BY CCT/TRIC | TUBB3 | TUBA1A, TUBB2B | MIRO1 |
| MICROTUBULE-DEPENDENT TRAFFICKING OF CONNEXONS FROM GOLGI TO THE PLASMA MEMBRANE | TUBB3 | TUBB2B, TUBA1A | MIRO1 |
| COPI-INDEPENDENT GOLGI-TO-ER RETROGRADE TRAFFIC | TUBB3 | TUBA1A, TUBB2B | MIRO1 |
| AGGREPHAGY | TUBB3 | TUBB2B, TUBA1A | MIRO1 |
| HCMV EARLY EVENTS | TUBB3 | TUBA1A, TUBB2B | MIRO1 |
| EML4 AND NUDC IN MITOTIC SPINDLE FORMATION | TUBB3 | TUBA1A, TUBB2B | MIRO1 |
| RESOLUTION OF SISTER CHROMATID COHESION | TUBB3 | TUBB2B, TUBA1A | MIRO1 |
| PKR-MEDIATED SIGNALING | TUBB3 | TUBA1A, TUBB2B | MIRO1 |
| MITOTIC PROMETAPHASE | TUBB3 | TUBB2B, TUBA1A | MIRO1 |
| COPI-MEDIATED ANTEROGRADE TRANSPORT | TUBB3 | TUBB2B, TUBA1A | MIRO1 |

| | | | |
|---|---|---|---|
| PREFOLDIN MEDIATED TRANSFER OF SUBSTRATE TO CCT/TRIC | TUBB3 | TUBB2B, TUBA1A, ACTB | MIRO1 |
| ASSEMBLY AND CELL SURFACE PRESENTATION OF NMDA RECEPTORS | TUBB3 | TUBA1A, TUBB2B | MIRO1 |
| POST-CHAPERONIN TUBULIN FOLDING PATHWAY | TUBB3 | TUBB2B, TUBA1A | MIRO1 |
| GAP JUNCTION ASSEMBLY | TUBB3 | TUBB2B, TUBA1A | MIRO1 |
| KINESINS | TUBB3 | TUBB2B, TUBA1A | MIRO1 |

**Table S6:** Drugs targeting genes involved in ROBO pathways.

| Drug | Targeted proteins | Experiments |
| --- | --- | --- |
| **4-PHENYLBUTYRIC ACID** | PRKCA | GBA |
| **ATALUREN** | RPS4X, RPL6, RPS27A, RPL13, RPL38, RPS7, RPL37A, RPL18A, RPS24, RPL27A | IPD, LRRK2 |
| **BEVACIZUMAB-AWWB** | CXCR4, PRKCA | SNCA, GBA |
| **BORTEZOMIB** | PSMD5 | SNCA |
| **CARFILZOMIB** | PSMD5 | SNCA |
| **CISPLATIN** | CXCR4 | SNCA |
| **DOCETAXEL ANHYDROUS** | RPL13 | LRRK2 |
| **HYDROCHLOROTHIAZIDE** | PRKCA | GBA |
| **INGENOL MEBUTATE** | PRKCA | GBA |
| **MIDOSTAURIN** | PRKCA | GBA |
| **PLERIXAFOR** | CXCR4 | SNCA |
| **QUERCETIN** | PRKCA | GBA |
| **RESVERATROL** | PRKCA | GBA |
| **THALIDOMIDE** | RPL13 | LRRK2 |
